# Modeling the Role of Platelet-Released Polyphosphates in Tissue-Factor-Initiated Coagulation under Flow

**DOI:** 10.64898/2026.03.19.713007

**Authors:** Sradha Ramesh Bhatt, Alexander G. Ginsberg, Stephanie A. Smith, James H. Morrissey, Aaron L Fogelson

## Abstract

**Background:** Activated platelets release polyphosphate (polyP), a linear polymer of inorganic phosphate residues, from dense granules. Experiments performed under no-flow conditions show that polyP alters the kinetics of tissue factor (TF) pathway reactions, accelerating FXI activation by thrombin and FV activation by FXa and thrombin, and may impact inhibition by tissue factor pathway inhibitor α (TFPIα). How polyP influences this pathway in conjunction with platelet deposition under flow remains understudied.

**Objectives:** To investigate how polyP-mediated acceleration of FV and FXI activation modulates thrombin generation under flow in TF-initiated coagulation.

**Methods:** We extended a previously validated mathematical model of platelet deposition and coagulation under flow to examine polyP-mediated effects following a small vascular injury during intravascular clotting. Simulations varied the surface density of TF exposed, wall shear rate, and plasma TFPIα concentration.

**Results:** PolyP shifts the threshold TF density for a thrombin burst to lower TF densities. For TF densities above this threshold, polyP shortens the lag time to thrombin generation in a TF- and shear-rate-dependent manner. Although no explicit effect of polyP on TFPIα function was included in the model, thrombin generation was much less sensitive to TFPIα concentration with polyP, in a TF-dependent manner. Relative contributions of accelerations of FV and FXI activations depend on incompletely known enhancements by polyP.

**Conclusions:** The experimentally observed influence of polyP on TFPIα function depends on TF density and may arise indirectly from accelerated FV and FXI activation, with the dominant effect arising through accelerated thrombin-mediated conversion of FV to FVa.

## Introduction

Platelet polyphosphate (polyP) has attracted growing attention as an important regulator of blood coagulation. It is a linear polymer of inorganic phosphate residues and is released from platelet dense granules upon platelet activation [1]. PolyP is ubiquitous across bacteria, fungi, plants, and animals [2]. It varies substantially in length with relatively short polymers (∼60-100 phosphate units) secreted by human platelets and much longer polymers (up to several thousand phosphate units) released by microorganisms. These length differences are functionally significant, as different polyP sizes differentially modulate the blood clotting system [3].

Studies show that polyP from both bacteria and human platelets plays important roles in inflammation and coagulation. Early clinical observations linked defects in platelet dense granules to bleeding tendencies [4,5]. Later studies provided a potential explanation for that link: defects in dense granules can lead to platelet polyP levels approximately a tenth of normal [6]. Moreover, persons with Hermansky-Pudlak syndrome have platelets lacking polyP and exhibit reduced clotting activity restored by the addition of platelet-sized polyP, highlighting the procoagulant role of platelet-sized polyP [7].

PolyP is highly anionic, binding tightly to proteins which initiate the contact pathway of coagulation. The first reported role for polyP in blood clotting demonstrated that polyP is strongly procoagulant and triggers the contact pathway [8]. Subsequent studies showed that long-chain polyP can potently activate the contact pathway and trigger thrombosis and inflammation [9].

PolyP also enhances the activation of key clotting factors in the tissue factor (TF) and common coagulation pathways. Studies have shown that platelet-released polyP accelerates the proteolytic activation of factor V (FV) by both factor Xa (FXa) and thrombin, resulting in an accelerated thrombin burst during plasma clotting reactions [8,10]. Platelet-sized polyP also is reported to decrease the activity of the coagulation inhibitor tissue factor pathway inhibitor α (TFPIα), possibly indirectly through its acceleration of FV activation [8]. Additionally, platelet-sized polyP accelerates factor XI (FXI) activation by thrombin (by ∼3000-fold), further supporting a role for polyP in amplifying thrombin generation [11,12].

Together, previous studies indicate that polyP is an important modulator of hemostasis and thrombosis. This has prompted increased interest in polyP as a therapeutic target [13], as inhibiting polyP function *in vivo* may provide a strategy for developing antithrombotic or anti-inflammatory agents, potentially with reduced bleeding risk compared with conventional anticoagulant therapies [14]. Also, since added polyP can shorten clotting times in plasma from patients with hemophilia A or B, platelet-sized polyP may be useful as a hemostatic agent [15]. In hemophilia plasma, the procoagulant effect of polyP was additive with recombinant activated factor VII, suggesting it could complement existing therapies [15].

To better understand the potential therapeutic roles of polyP, it is important to examine how polyP influences the coupled reactions of coagulation under flow. *In vitro* studies of polyP are generally performed under static, purified conditions, capturing key biochemical effects but overlooking the complexity of the interconnected reactions that regulate coagulation under flow. To this end, Yeon et al.[16] developed a mathematical model of the contact pathway of coagulation that examined the effect of surface-immobilized polyP localized on the walls of microfluidic channels during clot formation. Their simulations predicted that localized short-chain polyP can accelerate clotting of flowing blood plasma at sub-physiological shear rates. Their model treated all coagulation reactions as occurring in the moving plasma phase. However, it is known that critical coagulation reactions occur on the surfaces of activated platelets that have accumulated at the injury site, as included in our previous modeling studies [17,18]. Our studies indicated that without such surface binding, activated coagulation factors are rapidly cleared by flow, reducing their local concentrations and limiting thrombin generation. Motivated by these observations, we extend our existing, experimentally validated mathematical model of platelet deposition and TF-mediated coagulation under flow to investigate the role of polyP in modulating thrombin generation.

## Methods

Previous versions of the model on which this paper is based have been published [17–23] and their predictions experimentally validated [24–27]. Here, we sketch that model and indicate how we extend it to include platelet-released polyP.

### Review of our previous mathematical model of coagulation and platelet deposition

We consider a vessel the size of an arteriole or venule in which a small injury has uncovered a patch of subendothelium, exposing TF and collagen to flowing blood, triggering coagulation (Figure 1A) and platelet deposition (Figure 1B), respectively. We focus on events in a thin fluid layer above the injury, which we call the “reaction zone’’ (RZ) (Figure 1B) and whose initial height (∼1-2 µM) we estimated by a boundary layer analysis accounting for the near-wall flow velocity and the diffusivities of the relevant protein and cellular species [17].

**Figure 1:**
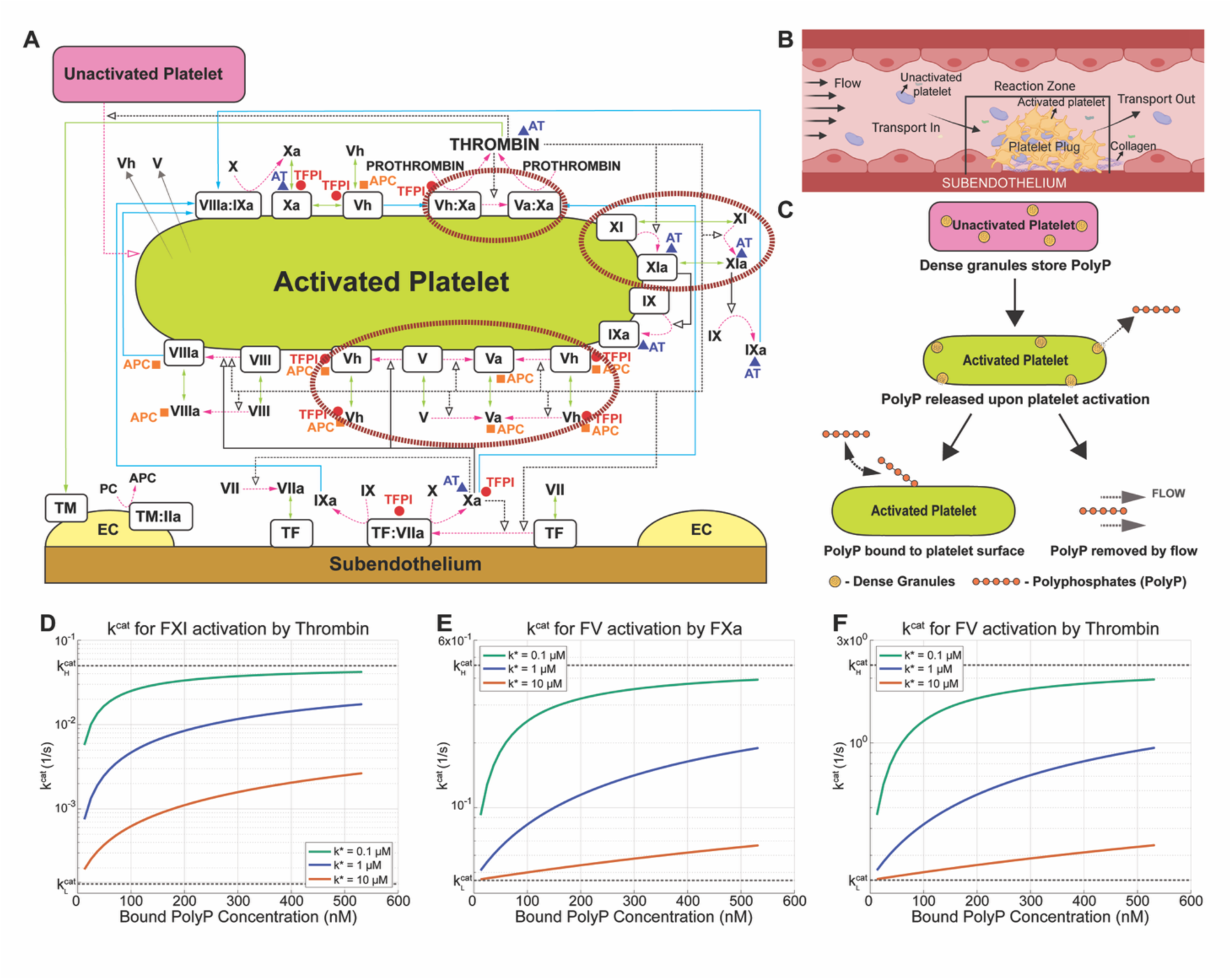
Schematic of coagulation reactions and representation of polyP dynamics. (A) The coagulation enzyme-cofactor complexes form on the subendothelium and APSs. Thrombomodulin (TM) on endothelial cells (ECs) is a cofactor for thrombin in producing the inhibitor activated protein C (APC). Other major inhibitors are antithrombin (AT) and TFPIα. Surface-bound enzyme complexes TF:VIIa, VIIIa:IXa, Va:Xa, and TM:IIa and other surface-bound species are shown in boxes. Also shown are: cell or chemical activation (*dashed magenta arrows*); movement in fluid or along a surface (*solid blue arrows*); enzyme action in a forward direction (*solid black arrows*); the feedback action of enzymes (*dashed black arrows*); binding to or unbinding from surface (*light green double-headed arrows*); chemical inhibitors (*red, orange and purple shapes*); and release of species from platelet stores, including FV and FVh (*grey arrows with faded tails*). The large circles highlight specific reactions that are accelerated by platelet-derived polyP in its presence. Adapted from [19]. (B) Vascular injury exposes collagen (*purple strands*). Platelets from upstream (*blue*) adhere to the collagen in the RZ, become activated (*spiky yellow*), and aggregate to form a platelet plug. Adapted from [23]. (C) PolyP released from activated platelets binds to APSs or is cleared by flow. (D-F) Dependence of the catalytic rate constant *k^cat^* on bound polyP concentration for three choices of *k\** and three coagulation reactions: (D) FXI activation by thrombin, (E) FV activation by FXa, and (F) FV activation by thrombin.

We consider all protein and cellular species within the RZ to vary over time but to be uniformly distributed in space and describe each species by its time-dependent concentration. We use an ordinary differential equation (ODE) to express the rate of change of each concentration due to reactions and transport into and out of the reaction zone. The overall model consists of coupled nonlinear ODEs for all species.

The model includes the number densities of three populations of platelets: unactivated fluid-phase platelets, activated platelets bound to the subendothelium, and activated platelets bound to other platelets in the thrombus. It includes mechanisms by which platelets become activated and bind to the subendothelium or other platelets along with mechanisms by which fluid-phase platelets move with the flow (Figure 1B). Bound platelets are considered immobile.

The model includes concentrations of zymogens, pro-cofactors, and the corresponding enzymes and active cofactors of the TF pathway (Figure 1A) and of the major chemical inhibitors of coagulation. It distinguishes clotting proteins in part by their chemical identity. Because the initiation of coagulation occurs on the subendothelial surface whereas critical reactions in the amplification phase of coagulation occur on activated platelet surfaces (APSs), the model further distinguishes proteins by whether they are in the plasma, bound to the subendothelium, or bound to an APS.

The model allows enzymes activated on the subendothelium to reach an APS by unbinding from the subendothelium, moving through the plasma, then binding to a receptor on the APS. While in the plasma, species may be carried downstream by the flow, removing them from the RZ.

Because APSs are the sites for critical procoagulant and inhibitory reactions, platelets play a prominent role in the model. Since each activated platelet has a limited ability to support these reactions, the increasing availability of APSs as platelets accumulate in the RZ strongly influences the progression of the coagulation reactions. Platelets also have an anticoagulant role. Namely, each platelet’s adhesion to the injured vessel wall covers part of the subendothelium and blocks access to TF:FVIIa on that portion, thus physically inhibiting the enzymatic activity of TF:FVIIa. Progressive increases in platelet adhesion block increasing fractions of the total TF:FVIIa activity.

We assume coagulation proteins interact as depicted in Figure 1 and as outlined here:

1. FVII and FVIIa bind to TF on the subendothelium. FXa activates FVII in plasma or in TF:FVII. FXa binds TF:FVII directly from the plasma without first binding to the subendothelium.
2. TF:FVIIa binds and activates FIX and FX from the plasma.
3. Thrombin in plasma or on an APS, and FXa on an APS, activate FV and FVIII. Prescribed numbers of unactivated and partially activated variants of FV are secreted from platelet α-granules upon platelet activation. We denote the partially activated form of FV as FVh (for half-activated) and the fully activated form as FVa [28]. FXa converts FV to FVh; thrombin converts FV and FVh to FVa.
4. Platelet-bound FVIIIa and FIXa bind to form the tenase complex FVIIIa:FIXa which activates platelet-bound FX, producing platelet-bound FXa.
5. Platelet-bound FVh and FVa bind platelet-bound FXa to form the prothrombinase complexes FVh:FXa (“PROh”) and FVa:FXa (“PRO”), respectively. These complexes can activate platelet-bound prothrombin into thrombin, which is immediately released into the plasma.
6. FXI is activated by thrombin in plasma and on an APS. FIX is activated by FXIa in plasma and on an APS.
7. The model includes the chemical inhibitors/inactivators antithrombin (AT), activated protein C (APC), and tissue factor pathway inhibitor (TFPIα).

- By binding to FIXa, FXa, FXIa, and thrombin, AT permanently inactivates them.
- APC binds to FVa and FVIIIa in the plasma and on an APS, permanently inactivating them. However, APC cannot bind to FVa or FVIIIa already incorporated in platelet-bound prothrombinase or tenase. APC is produced in the endothelial zone (EZ) adjacent to the RZ by a complex of thrombin and endothelial-cell-bound thrombomodulin. To bind to thrombomodulin, thrombin must diffuse from the RZ to the EZ; while, to affect coagulation dynamics, APC must diffuse from the EZ into the RZ.
- TFPIα in plasma binds to FXa. TFPIα: FXa can then inhibit TF:FVIIa. TFPIα also binds to and inhibits FXa and FVh in the plasma or on an APS, as well as to the FVh in PROh on an APS.
8. The activity of TF:FVIIa decreases as platelets adhere to the subendothelium, physically blocking access to the subendothelium patch to which they are attached.

### Model extensions to include effects of polyP

We explore the behavior of two variants of the above model with polyP. In the “implicit” model, we change only the rate constants for thrombin activation of FXI to FXIa, FV and FVh to FVa, and for FXa activation of FV to FVh, to reflect the acceleration of those reactions observed in the presence of platelet polyP. This is tantamount to assuming that a saturating concentration of polyP is present from the start of each simulation. In the implicit model we replace the low catalytic rate constant 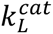 relevant when polyP is absent, with a high-rate constant 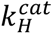 relevant when polyP is present. Default 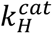 values are 10-fold and 300-fold the 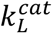 values for the polyP-enhanced FV and FXI activation reactions, respectively (see Table S1).

In the “explicit” model, we include the release of a specified quantity of polyP by each platelet as the platelet is activated. We assume that the released polyP polymers bind to specific sites on APSs or are washed away by the flow (Figure 1C). We assume that the effect of polyP on the coagulation reactions depends in a saturable way on the concentration of platelet-bound polyP (*P^b^*), as described by Equation (1) (see also Figures 1D-F),

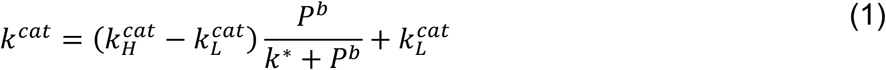

For the explicit model, we track the concentrations of fluid-phase polyP (*P^f^)* and platelet-bound polyP (*P*^b^). The total concentration of polyP (*P^f^* + *P*^b^) increases at the rate of polyP release by platelets and decreases at the rate of removal of fluid-phase polyP by flow (see Equations (S1-S2)). While bound to an APS, polyP is not susceptible to removal by flow. *P*^b^ increases as platelet-binding sites become more available, which occurs as platelets accumulate.

We assume that the kinetics of polyP binding to and unbinding from platelets are fast relative to the rates of release of polyP from platelet stores and of removal of polyP by flow. We therefore make the quasi-steady-state assumption that *P^f^*and *P^b^* equilibrate rapidly according to the current total polyP and polyP binding site concentrations (see Section S2). Consequently, the explicit model requires the addition of one differential equation for the total polyP concentration and algebraic formulas for *P^f^* and *P^b^*. Importantly, the model extension involves only three unknown parameters, the number *n_bs_* of binding sites per platelet for polyP polymers, the dissociation constant *K_D_* for polyP binding with these sites, and the concentration *k\** of bound polyP at which the catalytic rates for activation of FXI and FV species are halfway between their low and high values 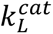 and 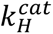. The model’s behavior for ranges of these parameters is examined.

## Results

Results from our simulations highlight changes, over the course of coagulation, in concentrations of proteins that polyP affects directly, FVh, FVa, FXIa, and others that polyP influences indirectly, including FVIIIa:FIXa, FVa:FXa, and thrombin. For thrombin we define two output metrics: lag time *t_lag_* and the thrombin concentration at 10 minutes [Thrombin]_10min_, which are widely used in the literature [29–31]. Lag time *t_lag_* is defined as the time at which [Thrombin] reaches 1 nM, a concentration at which thrombin activates platelets. [Thrombin]_10min_ is the thrombin concentration 10 minutes after TF exposure. We refer to curves of [Thrombin]_10min_ versus TF density as threshold curves. A typical threshold curve exhibits a small interval of TF density over which [Thrombin]_10min_ increases sharply from low levels (< 0.1 nM) with little physiological effect to high levels (>10 nM) with strong effect. We define the rapid increase in thrombin generation that occurs once the TF threshold is achieved as the *thrombin burst*.

### PolyP enhances thrombin generation, FXIa formation, and FVa formation

In Figure 2, we examine how varying TF density affects [Thrombin]_10min_ and *t_lag_* with and without polyP. We conducted a series of simulations at flow shear rates of 100, 500, and 1500 s^−1^, with [TFPIα]=0.5 nM, varying only TF density and computing the resulting thrombin concentration timecourse. Figure 2A shows that for shear rate 100 s^−1^ the addition of polyP decreases *t_lag_* for each TF density and can lead to a thrombin burst when none occurs without polyP (TF ∼0.05-0.4 fmol/cm^2^). Figure 2B shows that the threshold curves [Thrombin]_10min_ are shifted to the left when polyP is present. Even when a thrombin burst occurs without polyP, adding polyP increases [Thrombin]_10min_ by up to several orders of magnitude for low to moderate TF densities (TF ∼0.4-2 fmol/cm^2^). At high TF densities, however, polyP hardly affects [Thrombin]_10min_.

**Figure 2:**
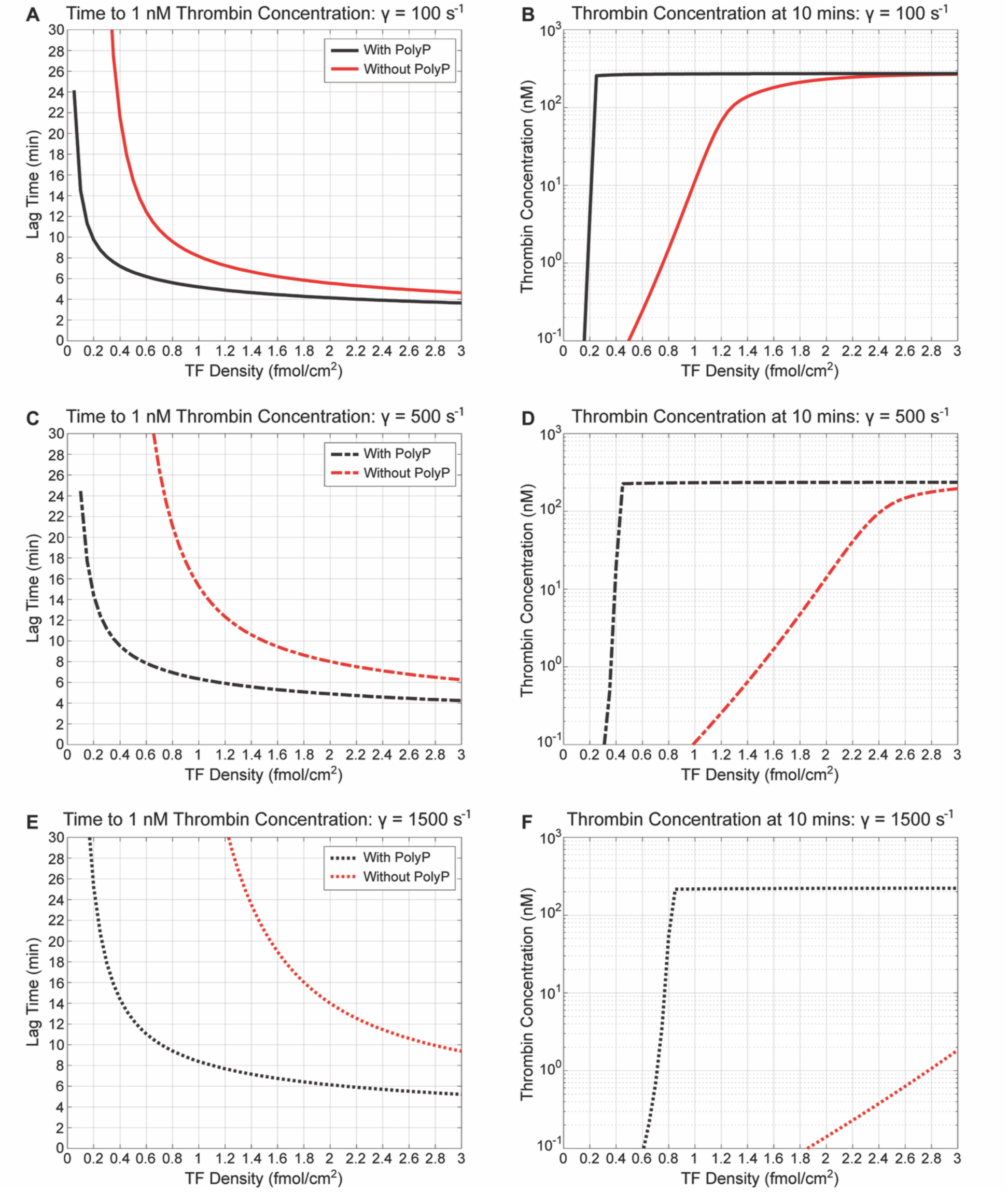
PolyP shortens the lag time and increases [Thrombin]_10min_ in the implicit model. Time to 1 nM thrombin (A, C, E) and **[Thrombin]_10min_** (B, D, F) with polyP (*black*) and without polyP (*red*) at [TFPIα] = 0.5 nM across shear rates (γ) of 100 s^−1^ (A-B, *solid lines*), 500 s^−1^ (C-D, *dash-dotted lines*), and 1500 s^−1^ (E-F, *dotted lines*).

PolyP’s effects on *t_lag_* and [Thrombin]_10min_ are consistent across shear rates as shown in Figures 2C-F. As shear rate increases, curves for lag time and the thrombin generation thresholds shift to the right. However, this shift is smaller in the presence of polyP than in its absence.

In Figure 3, we examine how the presence of polyP in the implicit model affects the time courses of thrombin, FXIa, and FVa concentrations at TF=1 fmol/cm^2^. Figure 3A shows that thrombin generation occurs sooner and increases more rapidly when polyP is present. To explain this, we focused on the direct effects of polyP in the implicit model, namely, acceleration of FXI activation by thrombin and acceleration of FV activation by thrombin and FXa. We examined the time courses of FXIa and FVa concentrations, along with their downstream effects.

**Figure 3:**
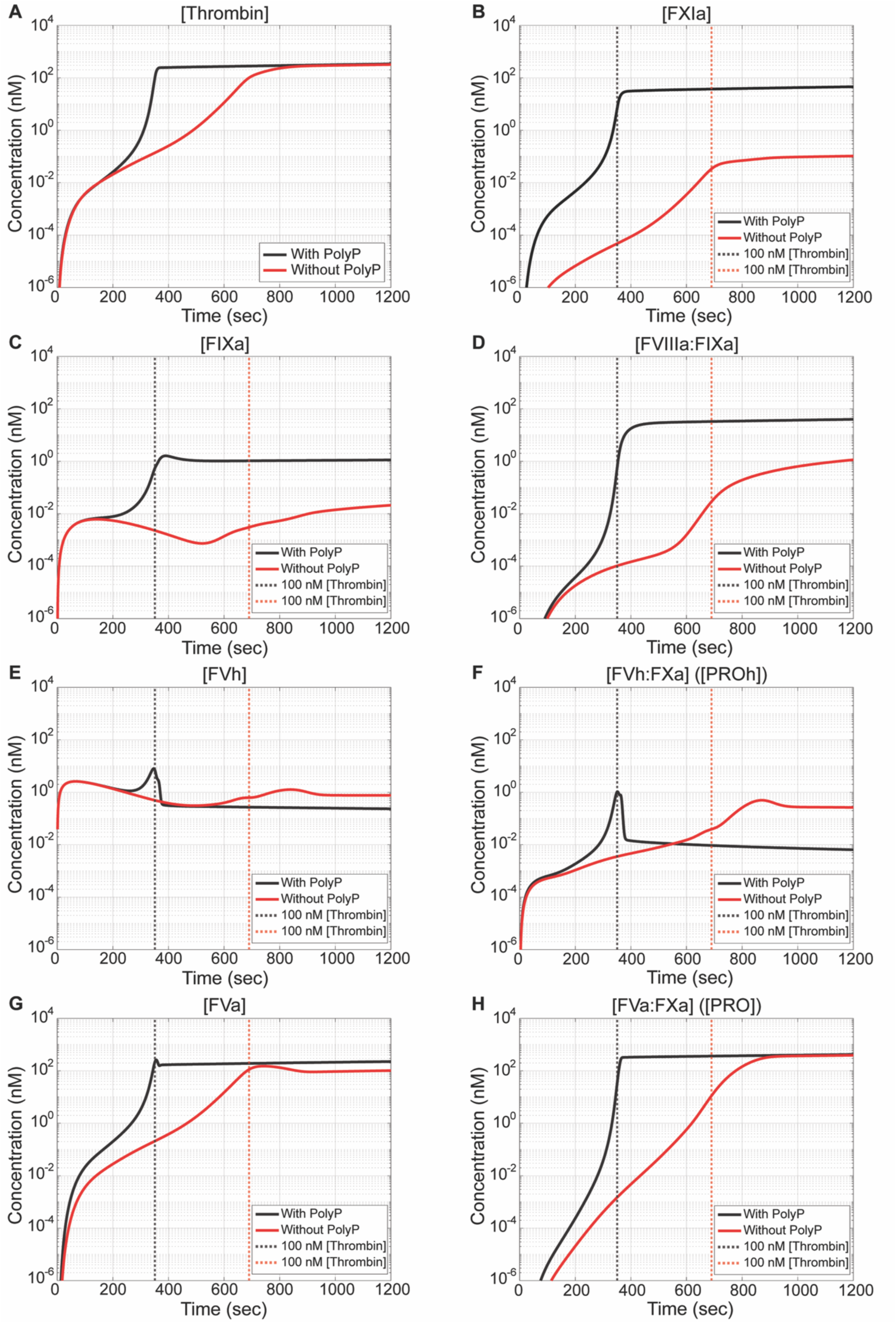
PolyP enhances the speed of thrombin generation in the implicit model. Time courses of enzyme concentrations with polyP (*black*) and without polyP (*red*): (A) Thrombin, (B) FXIa, (C) FIXa, (D) FVIIIa:FIXa (Tenase), (E) FVh, (F) FVh:FXa (PROh), (G) FVa, and (H) FVa:FXa (PRO). Vertical dashed lines indicate the time to reach 100 nM [thrombin] in each case. TF density = 1 fmol/cm^2^; [TFPIα] = 0.5 nM; shear rate = 100 s^−1^.

Figures 3B-D illustrate the consequences of accelerated FXI activation by thrombin when polyP is present. The FXIa concentration is dramatically enhanced and this, in turn, enhances the FIXa concentration through FXIa-dependent activation of FIX, and promotes FVIIIa:FIXa (tenase) complex formation on APSs. The combined effects of enhanced FXIa, FIXa and tenase concentrations with polyP contribute to the enhanced thrombin generation observed in Figure 3A.

Figures 3E-H illustrate the consequences of accelerated FV activation by FXa and thrombin when polyP is present. The accelerated reactions include conversion of FV to FVh by FXa, conversion of FV to FVa and FVh to FVa by thrombin, and conversion of PROh to PRO by thrombin’s cleavage of FVh within PROh to FVa within PRO. Figure 3E shows that polyP accelerates the formation of FVh, which exhibits a transient “bump.” The rise corresponds to the enhanced conversion of FV to FVh by FXa, while the decline occurs as FVh is incorporated into PROh and subsequently converted to FVa by thrombin. Figure 3F shows a similar transient bump for PROh. Here, the rise corresponds to increased binding of FVh to FXa to form PROh, and the decline occurs as thrombin converts PROh to PRO. In Figure 3G, FVa concentration increases earlier and reaches higher levels in the presence of polyP, which is driven by enhanced activation of FV and FVh by thrombin. The FVa concentration also exhibits a small transient bump similar to PROh in Figure 3F.

Figure 3H shows that polyP accelerates PRO formation, consistent with faster conversion of PROh to PRO and FV to FVa as thrombin concentration rises. PRO forms faster and earlier in the presence of polyP than in its absence. The direct enhancements of FVh, FVa, and PRO formation, along with indirect enhancement of PROh in the presence of polyP, together contribute to the faster thrombin generation observed in Figure 3A.

The results of Figure 3 collectively echo the experimentally observed roles of polyP in enhancing thrombin generation by accelerating FXI and FV activation [8,11]. We also examined the time courses of these species for TF density 0.3 fmol/cm^2^ (Figure S1). Although the interpretation is similar, we observed a large delay in thrombin generation without polyP, with the thrombin concentration reaching ∼100 nM only after 40 minutes as compared to ∼10 minutes when polyP is present.

### Relative contributions of polyP-accelerated FXI and FV activation to thrombin formation

In Figure 4, we show the extent to which polyP-accelerated FXI activation by thrombin and polyP-accelerated FV activation by thrombin and FXa act separately and in combination to promote thrombin generation. Figures 4A-B show that accelerating either FXI activation alone or FV activation alone decreases *t_lag_* (Figure 4A) and increases [Thrombin]_10min_ (Figure 4B). Each produces slightly more than half the effect of accelerating them together for TF densities between ∼0.05-0.1 fmol/cm^2^. Therefore, polyP-accelerated FXI and FV activation contribute roughly equally to accelerating thrombin generation in the implicit model, and their combined effect is generally sub-additive. At higher TF densities (>0.3 fmol/cm^2^), accelerated FV activation contributes slightly more than accelerated FXI activation.

**Figure 4:**
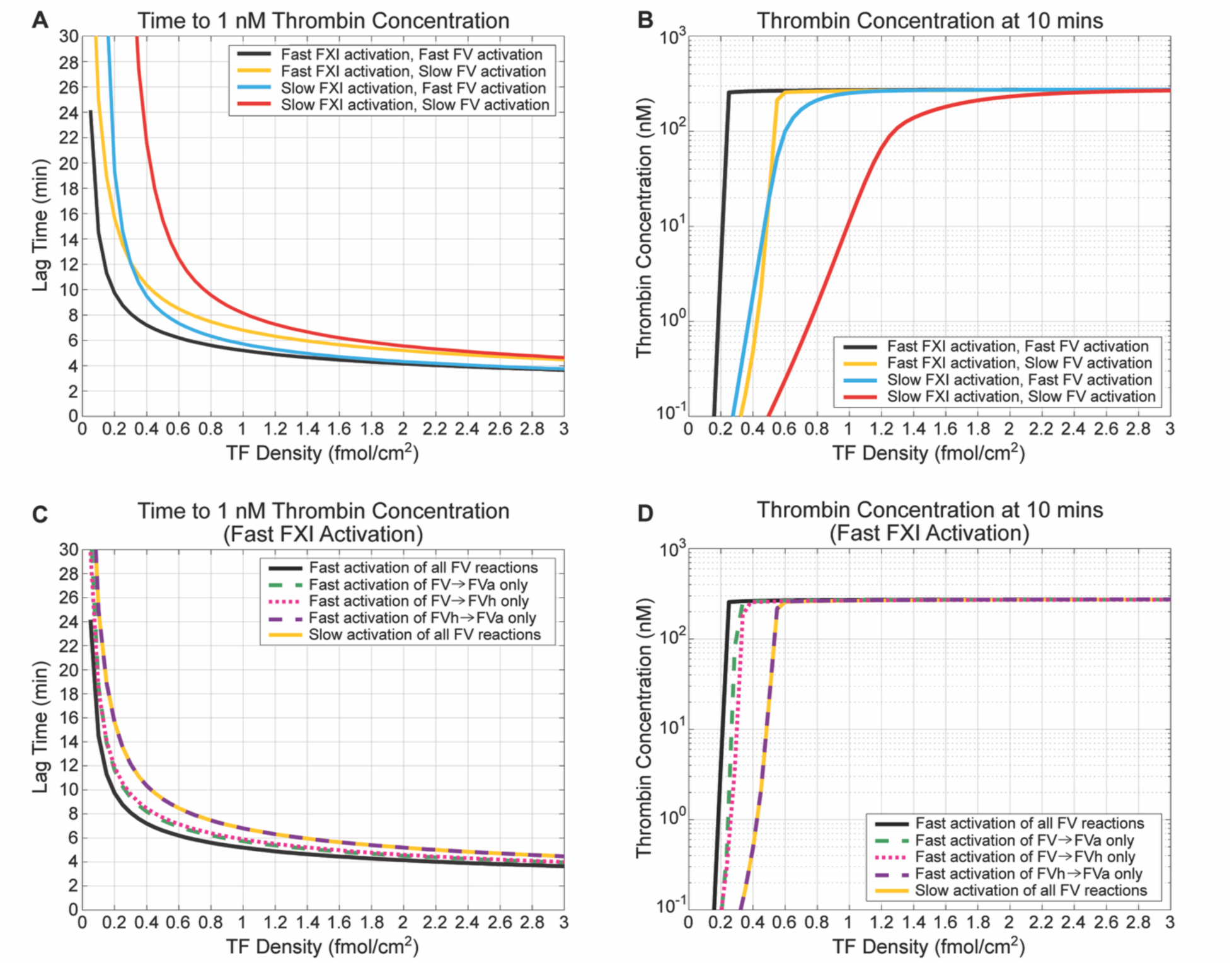
Relative contributions of FXI and FV activation to thrombin generation with polyP in the implicit model. (A) Time to 1 nM thrombin and (B) [thrombin] at 10 minutes with varying levels of acceleration of FXI activation by thrombin and FV activation by FXa and thrombin. (C) Time to 1 nM thrombin and (D) [Thrombin]_10min_ with accelerated FXI activation by thrombin and varying levels of acceleration of FV activation by FXa and thrombin. In (C) and (D), the curves corresponding to slow activation of all FV reactions (*solid yellow*) and to fast activation of FVh to FVa only (*dashed purple*) are on top of one another. “Fast” indicates that the respective reaction has been accelerated due to the presence of polyP, while “slow” indicates the baseline rate of activation without polyP. [TFPIα] = 0.5 nM; shear rate = 100 s^−1^.

Recall that polyP-accelerated FV activation has three parts: accelerated activation of FV to FVa and FVh to FVa by thrombin, and accelerated activation of FV to FVh by FXa. The accelerated conversion of FVh to FVa also includes conversion within PROh, yielding faster activation of PROh to PRO. Figures 4C-D show the effects of accelerating these different FV activation reactions. Accelerating either FV to FVa or FV to FVh alone decreases *t_lag_* across TF densities (Figure 4C) and increases [Thrombin]_10min_ (Figure 4D), although FV to FVa activation leads to slightly more thrombin formation, mainly for TF densities ∼0.2-0.6 fmol/cm^2^. Therefore, accelerated activation of FV to FVa and FV to FVh each contributes roughly equally to the acceleration of thrombin generation seen when polyP is included in the implicit model.

All simulations shown in Figure 4 include polyP-accelerated FXI activation by thrombin. In additional simulations (Figure S2) without this acceleration, faster activation of FV to FVa alone contributes more to the acceleration of thrombin generation than faster activation of FV to FVh alone. However, the effect of accelerating FVh to FVa activation by thrombin remains negligible.

Results obtained by repeating these simulations using a 100-fold increase in FV activation by FXa and thrombin, with and without accelerated FXI activation by thrombin, are shown in Figures S3, S4 and S5, respectively.

### PolyP reduces the sensitivity of thrombin generation to TFPIα concentration in a TF-dependent manner

In Figure 5, we examine polyP’s influence on TFPIα function. At TF=1 fmol/cm^2^, including polyP using the implicit model makes the system much less sensitive to the TFPIα concentration. With polyP (Figure 5A), *t_lag_* increases slowly from ∼4 to ∼9 minutes at 0 and 2.5 nM TFPIα, respectively. Without polyP, *t_lag_* rises quickly and steeply, from ∼6 to >30 minutes at 0 and 2.5 nM TFPIα, respectively. Figure 5B shows that [Thrombin]_10min_ is insensitive to TFPIα when polyP is present, remaining nearly constant and high (∼300 nM) across all TFPIα levels. In contrast, without polyP, [Thrombin]_10min_ drops by more than three orders of magnitude with increasing TFPIα (∼150 to 0.01 nM at 0 and 2.5 nM TFPIα, respectively). These results demonstrate that, although no explicit effect of polyP on TFPIα function was included in the simulations, *t_lag_* and [Thrombin]_10min_ were much less sensitive to TFPIα concentration with polyP-accelerated FXI and FV activation than without them. Similar behavior is observed at TF density 0.3 fmol/cm^2^ (Figure S6).

**Figure 5:**
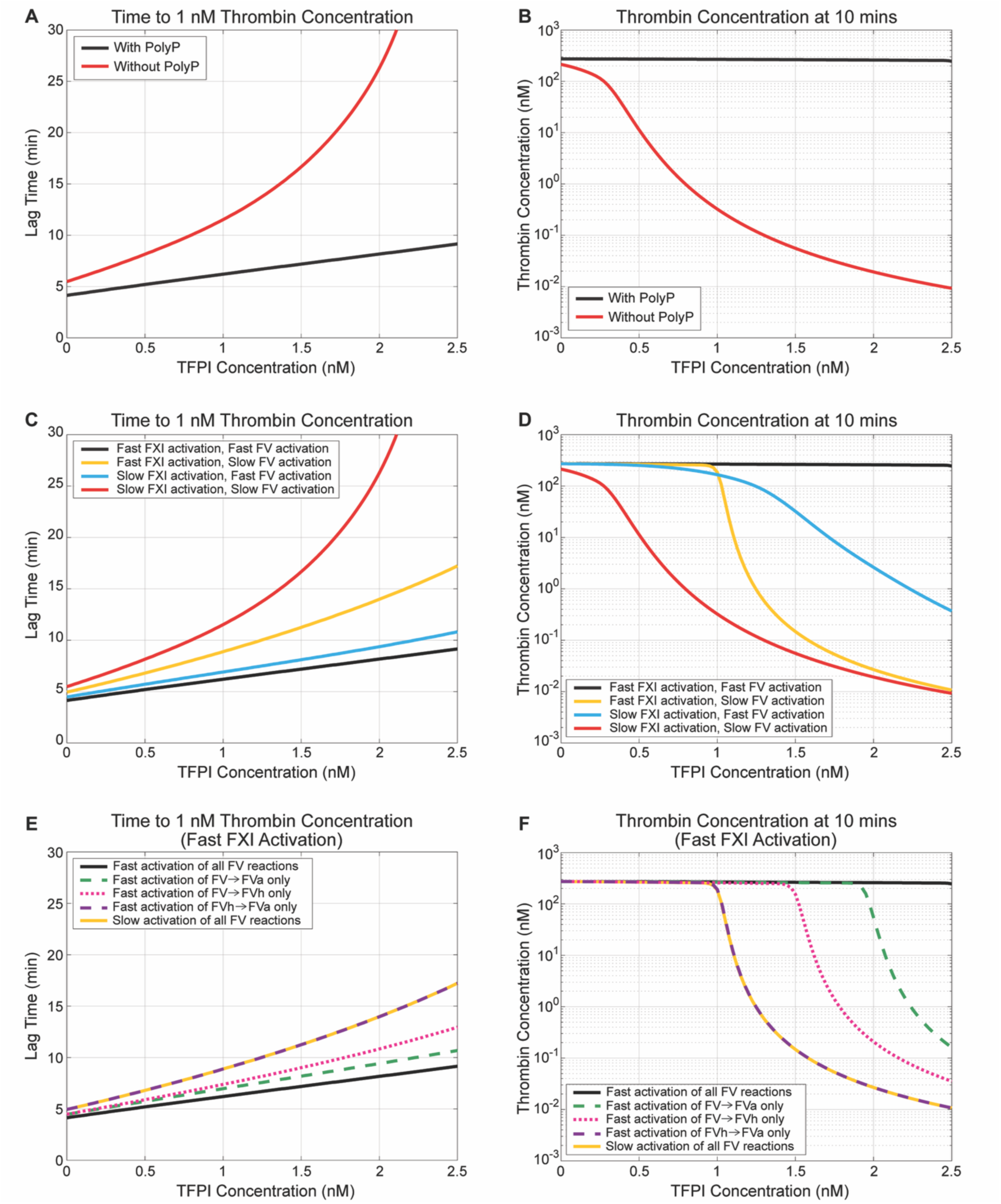
PolyP diminishes the sensitivity of thrombin generation to TFPI function in the implicit model. (A-F) Time to 1 nM thrombin (*left*) and [Thrombin]_10min_ (*right*) versus TFPIα concentration: (A-B) with polyP (*black*) and without polyP (*red*), (C-D) with varying levels of acceleration of FXI activation by thrombin and FV activation by FXa and thrombin, and (E-F) with accelerated FXI activation by thrombin and varying levels of acceleration of FV activation by FXa and thrombin. In (E-F), the curves corresponding to slow activation of all FV reactions (*solid yellow*) and fast activation of FVh to FVa only (*dashed purple*) overlap. “Fast” indicates that the respective reaction is accelerated due to the presence of polyP, while “slow” indicates the baseline rate of activation without polyP. TF density = 1 fmol/cm^2^; shear rate = 100 s^−1^.

Figures 5C-D examine the relative contributions of polyP-accelerated FXI and FV activation to TFPIα insensitivity. Figure 5C shows that accelerating either FXI or FV activation reduces the sensitivity of *t_lag_* to TFPIα. When only FV activation is accelerated, *t_lag_* remains short across TFPIα concentrations, increasing from ∼4 to ∼11 minutes at 0 and 2.5 nM TFPIα, respectively. When only FXI activation is accelerated, *t_lag_* rises more quickly with TFPIα, from ∼5 to ∼17 minutes at 0 and 2.5 nM TFPIα, respectively. Therefore, accelerating FV activation alone reduces the sensitivity of *t_lag_* to TFPIα more than does accelerating FXI activation alone.

Figure 5D shows the effects of varying the rates of FXI and FV activation on [Thrombin]_10min_. Accelerating either FXI or FV activation substantially reduces the sensitivity of [Thrombin]_10min_ to TFPIα at lower TFPIα concentrations (0-1 nM), with thrombin levels remaining nearly constant and high (∼200-300 nM) up to 1 nM TFPIα. When FV activation alone is accelerated, [Thrombin]_10min_ decreases gradually at higher TFPIα concentrations, from ∼200 to ∼0.4 nM at 1 and 2.5 nM TFPIα, respectively. In contrast, when FXI activation alone is accelerated, thrombin levels decline more sharply beyond 1 nM TFPIα, from ∼200 to ∼0.01 nM at 1 and 2.5 nM TFPIα, respectively. Thus, acceleration of FV activation alone reduces the sensitivity of [Thrombin]_10min_ to TFPIα more than does acceleration of FXI activation alone, consistent with the results for *t_lag_*. For both metrics, the least sensitivity to TFPIα is observed when both FXI and FV activation are accelerated. Moreover, [Thrombin]_10min_, remains high across all TFPIα concentrations. Taken together, Figures 5C-D show that while acceleration of both FXI and FV activation contributes to reduced TFPIα sensitivity, acceleration of FV activation has the larger overall effect on both thrombin metrics.

Figures 5E-F show the effects of accelerating the three types of FV activation reactions included in the model, with fast FXI activation. They show that accelerating either thrombin’s conversion of FV to FVa or FXa’s conversion of FV to FVh decreases the sensitivity of both [Thrombin]_10min_ and *t_lag_* to TFPIα, with the former producing the larger reduction. However, accelerating thrombin’s conversion of FVh to FVa alone had almost no effect on either metric.

The simulations in Figures 5E-F include polyP-accelerated activation of FXI by thrombin. Figure S7 shows additional simulations without this acceleration. Figure S8 shows results obtained by repeating the simulations in Figure 5 with a 100-fold increase in FV activation by FXa and thrombin along with accelerated FXI activation by thrombin.

The sensitivity of the thrombin generation metrics *t_lag_* and [Thrombin]_10min_ to TFPIα, described above for TF density 1 fmol/cm^2^, is decreased when polyP is present for higher TF densities (not shown). Figure 6 demonstrates that the sensitivity of thrombin generation to TFPIα depends both on the presence of polyP and the TF density. Figures 6A-B show that with polyP, both thrombin generation metrics become increasingly sensitive to TFPIα as the TF density is decreased below 1 fmol/cm^2^, with the greatest changes in sensitivity occurring for TF less than 0.5 fmol/cm^2^. Figures 6C-D show that without polyP, the thrombin metrics become increasingly insensitive to TFPIα as TF is increased from 1 to 3 fmol/cm^2^, although even for TF 3 fmol/cm^2.^ the metrics without polyP are more sensitive to TFPIα than they are for TF 1 fmol/cm^2^ with polyP.

**Figure 6:**
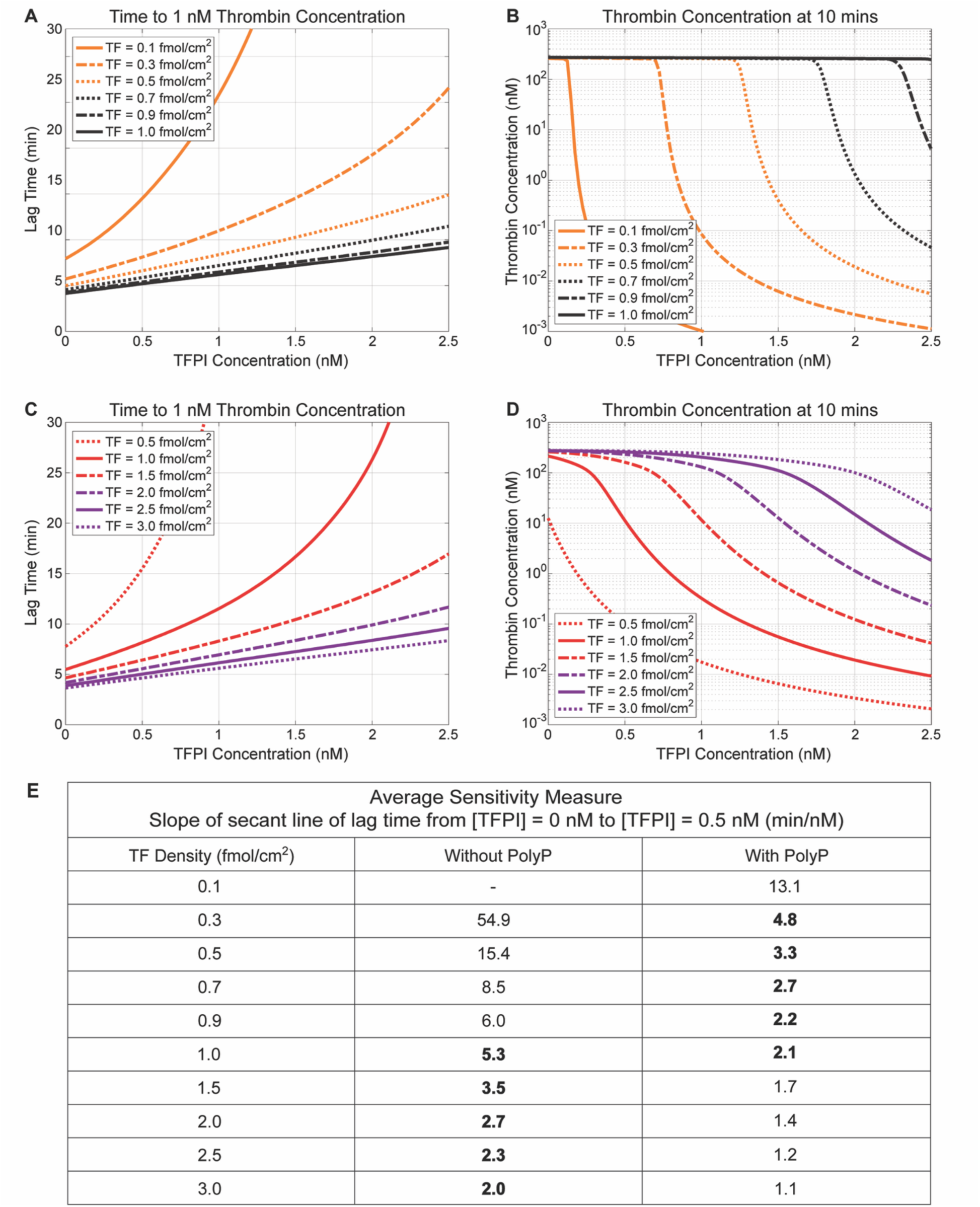
PolyP modulates the sensitivity of thrombin generation to TFPIα in a TF-dependent manner. (A-D) Time to 1 nM thrombin (*left*) and [Thrombin]_10min_ (*right*) versus TFPIα concentration: (A, B) with polyP for TF densities 0.1, 0.3, 0.5, 0.7, 0.9, and 1 fmol/cm^2^, and (C, D) without polyP for TF densities 0.5, 1, 1.5, 2, 2.5, and 3 fmol/cm^2^. (E) Average sensitivity measure (slope of secant line of lag time from 0 to 0.5 nM TFPIα) with and without polyP for the same TF densities as in (A-D). Bold entries indicate similar sensitivity slopes, occurring at lower TF with polyP and higher TF without polyP. The sensitivity jump refers to the difference in the sensitivity measure (slope of the secant line) between TF densities corresponding to the bold entries. This difference is smaller at low TF values with polyP and larger at higher TF values without polyP. A “-” in the table indicates that [thrombin] for the corresponding TF density never reached 1 nM for some TFPIα concentrations between 0 and 0.5 nM. Shear rate = 100 s^−1^.

The table in Figure 6E shows the sensitivity of the model’s behavior both with and without polyP in terms of the slope of the secant line of *t_lag_* versus TFPIα concentration from 0 to 0.5 nM. It emphasizes that while the inclusion of polyP makes thrombin production less sensitive to TFPIα for particular TF densities, thrombin production with polyP included is just as sensitive to TFPIα at lower TF densities as thrombin production without polyP is at higher TF densities. However, for each TF density, the slope of the secant line of *t_lag_* with polyP is smaller than the corresponding slope without polyP, indicating reduced sensitivity of thrombin generation to TFPIα in the presence of polyP. This is consistent with the results observed in Figure 5 for 1 fmol/cm^2^ TF. These results are independent of the choice of secant line (Figure S9).

### Explicit model parameters determine the impact of platelet-released polyP

Recall that the explicit model tracks polyP secretion from platelets and binding to activated platelet surfaces (APSs), and that catalytic rates depend dynamically on the concentration of bound polyP. The key parameters are the number of binding sites (*n_bs_*) for polyP on each APS, the dissociation constant (*K_D_*) for polyP binding to these sites, and the sensitivity (*k\**) of the catalytic rate constants to the bound polyP concentration (Equation 1). To our knowledge, these parameters have not been measured. We therefore vary these parameters widely to assess their impact. These parameters’ influence on the thrombin metrics *t_lag_* and [Thrombin]_10_ are shown in Figures 7A and 7B, respectively. Varying the explicit model parameters shifts both metrics between the extreme cases observed in the implicit model with and without polyP.

**Figure 7:**
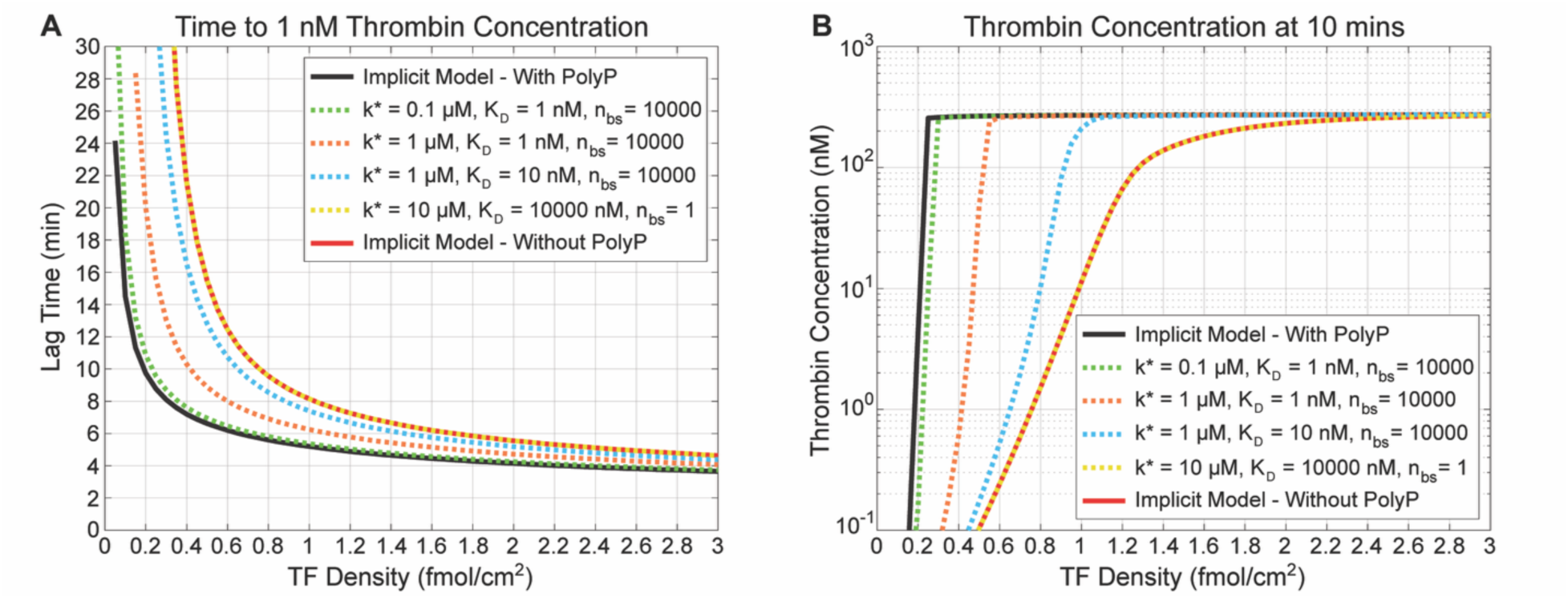
Explicit model parameters determine whether thrombin generation resembles the implicit model with or without polyP. Time to 1 nM thrombin (A) and [Thrombin]_10min_ (B) for the implicit model with polyP (*black*), the implicit model without polyP (*red*), and different explicit model parameter sets (*dotted*). In (A) and (B), the curves corresponding to the explicit model parameter set, *k\** = 10 uM, *K_D_* = 10000 nM and *n_bs_* = 1 (*dotted yellow*), and to the implicit model with polyP (*solid red*), are on top of one another. [TFPIα] = 0.5 nM; shear rate = 100 s^−1^.

The parameters’ effects on *t_lag_* and [Thrombin]_10_ are TF dependent. For low TF (0.7 fmol/cm^2^) lag times from 6 to 15 min are seen while for TF 1.5 fmol/cm^2^, a much smaller range, 5 to 9 min, is observed depending on the polyP kinetic parameters. Consequently, [Thrombin]_10min_ also varies strongly with these parameters and TF levels, ranging between 0.5 - 250 nM for 0.7 fmol/cm^2^ TF and 100 - 300 nM for 1.5 fmol/cm^2^ TF.

In the explicit model, the extent of acceleration of thrombin generation is captured roughly in the parameter ratio R = (*n_bs_*/*K_D_*)/*k\** (nM^−2^); it is directly proportional to the number *n_bs_* of polyP binding sites, and inversely proportional to both the dissociation constant *K_D_* and the sensitivity parameter *k\** in the catalytic rate expression for FXI and FV activation. In Figure 8, we examine nine parameter sets for which R is 0.1, 1, or 10 nM^−2^ (three sets for each R value). For each TF density, parameter sets with the same ratio yield nearly identical *t_lag_* and [Thrombin]_10min_. Increasing R accelerates thrombin production. For R=0.1, *t_lag_* decreases and [Thrombin]_10min_ increases only slightly compared to the case without polyP. However, increasing R to 1 and then to 10 strongly reduces *t_lag_* and strongly increases [Thrombin]_10min_, reaching about half the acceleration achieved in the implicit model. For even larger values of R, such as R=100 (not shown), *t_lag_* and [Thrombin]_10min_ are close to those of the implicit model, suggesting that the ratio R captures the dependence of thrombin metrics on the explicit model parameters. Figure S10 shows the time courses of thrombin, fluid-phase polyP and platelet-bound polyP for R=0.1, 1, or 10 nM^−2^.

**Figure 8:**
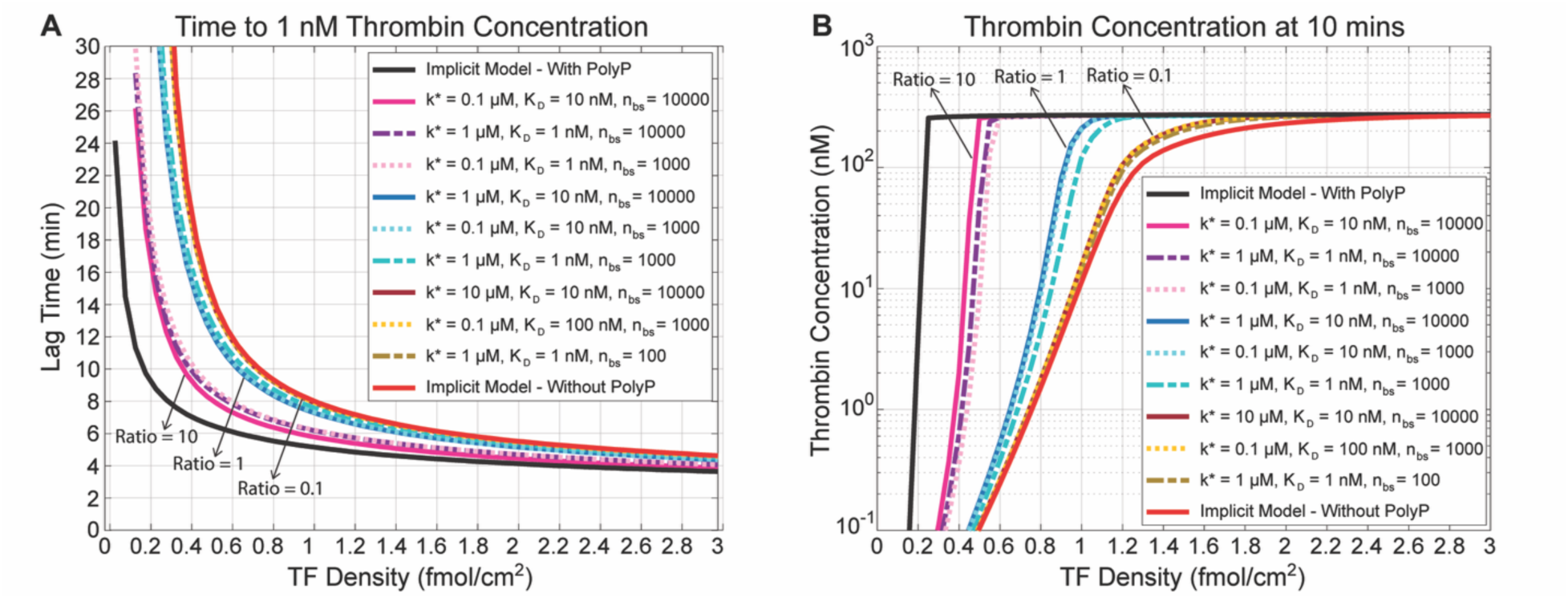
Effect of the explicit model parameter ratio, R, on thrombin generation. (A) Time to 1 nM thrombin *t_lag_* and (B) [Thrombin]_10min_ for the implicit model with polyP (*black*) and without polyP (*red*), and for explicit model parameter sets chosen to yield different values of the ratio R = (*n_bs_*/*K_D_*)/*k\** (nM^−2^). In (A), the red curve overlaps the brown curves. [TFPIα] = 0.5 nM; shear rate = 100 s^−1^.

## Discussion

To gain quantitative insight into the role of polyP in thrombin production, we extended an existing, validated mathematical model of flow-mediated platelet deposition and TF-initiated coagulation. Experiments show that polyP accelerates specific coagulation reactions, including FXI and FV activation [8,11]. Our modeling framework allows these effects to be examined within a full model of the TF pathway under flow that accounts for concurrent platelet accumulation and for a wider range of reaction conditions than in experiments, which are often performed under no-flow conditions and examine only a subset of coagulation reactions.

Using the model, we quantified the relative contributions of faster FXI and FV activation to thrombin formation. When polyP accelerates FV activation 10-fold, polyP-accelerated FXI and FV activation individually contribute approximately equally to accelerating thrombin generation, while their combined effect is sub-additive. Moreover, when FXI activation is enhanced, accelerated conversion of FV to FVa by thrombin and FV to FVh by FXa contribute roughly equally to the acceleration of thrombin generation, whereas accelerated conversion of FV to FVh by thrombin has a negligible impact. The last occurs because PROh and PRO activate prothrombin with the same kinetics [32].

Experiments in the presence of platelet-sized polyP have also shown apparent insensitivity to TFPIα levels but could not distinguish whether this is due to direct inhibition of TFPIα or is an indirect effect of accelerated coagulation reactions [8]. In our simulations, polyP indeed greatly decreases the sensitivity of thrombin generation to TFPIα. This change is driven by faster FV and FXI activation, with faster thrombin-mediated conversion of FV to FVa being the dominant contribution when FV activation is accelerated 10-fold. We emphasize that the model includes no direct action of polyP on TFPIα. The model further reveals that polyP’s ability to reduce TFPIα’s anticoagulant effect depends strongly on TF density. While polyP makes thrombin generation less sensitive to TFPIα at each TF density, it also makes thrombin production at lower TF densities about as sensitive to TFPIα as thrombin production without polyP is at higher TF densities. Thus, polyP non-uniformly affects TFPIα’s apparent activity in a TF-dependent manner.

Because quantitative measurements of polyP-enhanced FV activation by thrombin and FXa have not been reported [8,10], we assumed 10-fold increases in their catalytic rates (100-fold also tested; see Supplementary Material). Although the exact acceleration is unknown, our choice reproduces the experimentally observed reduction in lag time to 10 nM thrombin in the presence of polyP [8], validating our choice. However, simulations using a 100-fold increase in FV activation by FXa and thrombin alter TFPIα insensitivity and the relative importance of accelerated FV and FXI activations, motivating future experimental studies to quantify how much polyP accelerates FV activation.

In our explicit model, the impact of platelet-released polyP on thrombin generation depends on the parameter ratio R = (*n_bs_*/*K_D_*)/*k\** (nM^−2^). R increases with the number of polyP binding sites (*n_bs_*) on platelets and decreases with the dissociation constant (*K_D_*) for polyP binding and the sensitivity parameter (*k\**) in the catalytic rate for FXI and FV activation. Because the value of R influences thrombin generation and polyP concentrations, experimentally measuring the parameters that comprise this ratio is warranted although outside the scope of this study.

Future model explorations could incorporate additional experimentally reported effects of polyP whose quantitative details are unknown. These include FV activation by FXIa, likely most relevant for the contact pathway, polyP-enhanced FXI autoactivation, and FXIa-mediated TFPIα inactivation [10,11,33].

Our mathematical modeling results provide a coherent interpretation of how platelet-released polyP modulates thrombin generation under flow in TF-mediated coagulation. By accelerating FXIa and FVa formation, PolyP promotes earlier and stronger thrombin generation. PolyP indirectly reduces the sensitivity of thrombin generation to TFPIα in a TF-dependent manner, an effect predominantly due to polyP-accelerated thrombin-mediated activation of FV. This work provides mechanistic insight into how polyP may be exploited as either a hemostatic agent or an antithrombotic target.

## Author contributions

<u>S.Ramesh Bhatt, A.Ginsberg, and A.Fogelson</u> designed the model. S.Ramesh Bhatt, <u>A.Ginsberg, and A.Fogelson</u> implemented the model. <u>S.Ramesh Bhatt performed simulations. S.Ramesh Bhatt, A.Ginsberg, S.Smith, J.Morrissey, and A.Fogelson</u> analyzed and interpreted simulations. <u>S.Ramesh Bhatt, A.Ginsberg, and A.Fogelson wrote the</u> original paper draft. <u>S.Ramesh Bhatt, A.Ginsberg, S.Smith, J.Morrissey, and A.Fogelson</u> edited paper drafts.

