## Supplemental Material for "Modeling the Role of Platelet-Released Polyphosphates in Tissue-Factor-Initiated Coagulation under Flow"

Sradha Ramesh Bhatt[1], Alexander G. Ginsberg[1], Stephanie A. Smith[2], James H. Morrissey[2], and Aaron L. Fogelson[1, 3]
[1]Department of Mathematics, University of Utah, Salt Lake City, UT, USA
[2]Department of Biological Chemistry, University of Michigan Medical School, Ann Arbor, Michigan, USA
[3]Department of Biomedical Engineering, University of Utah, Salt Lake City, UT, USA

#### Supplemental Material

### 1 Explicit Model Formulation

This section describes the complete formulation of the explicit model for polyP dynamics. Our simulations do not directly use this representation; instead, a quasi-steady-state approximation obtained from this formulation is used (see Section 2). The explicit model tracks two polyP species: plasma-phase polyP ($P^{\mathrm{f}}$) and bound polyP ($P^{\mathrm{b}}$). The total polyP concentration ($P^{\mathrm{tot}}$) is defined as their sum. The corresponding ODEs are:

$$\frac{dP^{\mathrm{f}}}{dt} = \underbrace{n_{\mathrm{rel}}\frac{d}{dt}(P_S + P_V)}_{\substack{\text{release of polyP}\\ \text{upon platelet activation}}} - \underbrace{k_{\mathrm{flow}}P^{\mathrm{f}}}_{\substack{\text{flow-mediated}\\ \text{removal of polyP}}} - \underbrace{k_+\left\{n_{\mathrm{bs}}(P_S+P_V) - m_{\mathrm{bs}}P^{\mathrm{b}}\right\}P^{\mathrm{f}}}_{\text{binding of polyP to APSs}} + \underbrace{k_-P^{\mathrm{b}}}_{\substack{\text{unbinding of polyP}\\ \text{from APSs}}}, \tag{1}$$

$$\frac{dP^{\mathrm{b}}}{dt} = \underbrace{k_+\left\{n_{\mathrm{bs}}(P_S+P_V) - m_{\mathrm{bs}}P^{\mathrm{b}}\right\}P^{\mathrm{f}}}_{\text{binding of polyP to APSs}} - \underbrace{k_-P^{\mathrm{b}}}_{\substack{\text{unbinding of polyP}\\ \text{from APSs}}}, \tag{2}$$

where $n_{\mathrm{rel}}$ is the number of polyP polymers released by a platelet upon activation, $P_S$ and $P_V$ denote the concentrations of activated platelets that are attached and not attached to the subendothelium, respectively; $k_{\mathrm{flow}}$ is a rate function describing the flow-mediated removal of polyP [1]; $k_+$ and $k_-$ are the binding and unbinding rate constants for polyP polymers to and from activated platelet surfaces (APSs) respectively; $n_{\mathrm{bs}}$ is the number of polyP polymers that can bind to an APS; and $m_{\mathrm{bs}}$ is the number of binding sites occupied by a single polyP polymer (set to 1 in the model). The first term in Eq. (1) represents the rate of polyP release from APSs, while the second term describes its removal by flow. The third term accounts for the rate of polyP binding to APSs, where $n_{\mathrm{bs}}(P_S + P_V) - m_{\mathrm{bs}}P^b$ gives the number of available binding sites for polyP polymers on APSs. The final term gives the rate at which polyP unbinds from APSs. Similarly, the first term in Eq. (2) represents the rate of polyP binding to APSs and the second term corresponds to the rate at which polyP unbinds from APSs.

### 2 Quasi-Steady-State Approximation

In the quasi-steady-state approximation, the total polyP concentration ($P^{\mathrm{tot}}$) is tracked explicitely via an ODE, while the bound and free polyP concentrations, $P^{\mathrm{b}}$ and $P^{\mathrm{f}}$, are determined from via an algebraic equation. We assume that polyP released by activated platelets binds and unbinds rapidly to and from APSs, treating these reactions as fast processes compared to polyP release and platelet activation. $P^{\mathrm{b}}$ is the fast variable that quickly reaches a quasi-steady-state. To capture the separation of timescales for the slow-fast dynamics, we non-dimensionalize time as $\tau = k_{\mathrm{flow}}t$, giving $\frac{d}{dt} = k_{\mathrm{flow}}\frac{d}{d\tau}$. Then Eq. (2) becomes

$$k_{\mathrm{flow}}\frac{dP^{\mathrm{b}}}{d\tau} = k_+\left\{n_{\mathrm{bs}}(P_S+P_V) - m_{\mathrm{bs}}P^{\mathrm{b}}\right\}P^{\mathrm{f}} - k_-P^{\mathrm{b}}.$$

Dividing through by $k_+$, defining $\varepsilon = k_{\text{flow}}/k_+ \ll 1$, $K_D = k_-/k_+$, and setting $m_{\text{bs}} = 1$, gives

$$\varepsilon \frac{dP^{\text{b}}}{d\tau} = \left\{ n_{\text{bs}}(P_S + P_V) - P^{\text{b}} \right\} P^{\text{f}} - K_D P^{\text{b}}.$$

In the quasi-steady-state limit ($\varepsilon \to 0$), the left-hand side vanishes, yielding

$$0 = \left\{ n_{\text{bs}}(P_S + P_V) - P^{\text{b}} \right\} P^{\text{f}} - K_D P^{\text{b}}.$$

Solving gives the steady-state bound polyP concentration,

$$P^{\text{b}}_{\text{ss}} = \frac{n_{\text{bs}}(P_S + P_V)P^{\text{f}}}{K_D + P^{\text{f}}}. \tag{3}$$

Let $P^{\text{tot}} = P^{\text{f}} + P^{\text{b}}_{\text{ss}}$. Then,

$$P^{\text{tot}} = P^{\text{f}} + \frac{n_{\text{bs}}(P_S + P_V)P^{\text{f}}}{K_D + P^{\text{f}}}. \tag{4}$$

The ODE for total polyP concentration is obtained by summing Eq. (1) and Eq. (2) as

$$\frac{dP^{\text{tot}}}{dt} = n_{\text{rel}} \frac{d}{dt}(P_S + P_V) - k_{\text{flow}} P^{\text{f}}, \tag{5}$$

Rearranging (4) produces a quadratic equation in $P^{\text{f}}$. Taking the positive root and substituting into (5) gives the following ODE for $P^{\text{tot}}$:

$$\frac{dP^{\text{tot}}}{dt} = n_{\text{rel}} \frac{d}{dt}(P_S + P_V) - k_{\text{flow}} \left[ \frac{-n_{\text{bs}}(P_S + P_V) - K_D + P^{\text{tot}}}{2} + \frac{1}{2} \sqrt{\begin{array}{l} (n_{\text{bs}}(P_S + P_V))^2 + 2n_{\text{bs}}(P_S + P_V)K_D \\ - 2n_{\text{bs}}(P_S + P_V)P^{\text{tot}} + K_D^2 + 2K_D P^{\text{tot}} + P^{\text{tot}\,2} \end{array}} \right]. \tag{6}$$

In our simulations, the ODE for $P^{\text{tot}}$ is solved numerically. Substituting $P^{\text{f}} = P^{\text{tot}} - P^{\text{b}}_{\text{ss}}$ into Eq. (3), the resulting quadratic in $P^{\text{b}}_{\text{ss}}$ can be solved for the positive root, yielding

$$P^{\text{b}}_{\text{ss}} = \frac{n_{\text{bs}}(P_S + P_V) + K_D + P^{\text{tot}}}{2} - \frac{1}{2} \sqrt{\begin{array}{l} (n_{\text{bs}}(P_S + P_V))^2 + 2n_{\text{bs}}(P_S + P_V)K_D \\ - 2n_{\text{bs}}(P_S + P_V)P^{\text{tot}} + K_D^2 + 2K_D P^{\text{tot}} + P^{\text{tot}\,2} \end{array}}. \tag{7}$$

**Table S1:** Implicit and Explicit model parameters and kinetic constants.

| Parameter | Description | Value (Units) | Note |
|---|---|---|---|
| $k^{\text{cat}}_H$ | FXI → FXIa by thrombin | $0.05\ \text{s}^{-1}$ | [2] |
| | FV → FVa, FVh → FVa by thrombin | $2.3\ \text{s}^{-1}$ | Default |
| | FV → FVh by FXa | $0.46\ \text{s}^{-1}$ | Default |
| $k^{\text{cat}}_L$ | FXI → FXIa by thrombin | $1.29 \times 10^{-4}\ \text{s}^{-1}$ | [3] |
| | FV → FVa, FVh → FVa by thrombin | $0.23\ \text{s}^{-1}$ | [4] |
| | FV → FVh by FXa | $0.046\ \text{s}^{-1}$ | [5] |
| $n_{\text{rel}}$ | PolyP polymers released per activated platelet | $6.0 \times 10^4$ | [6] |
| $n_{\text{bs}}$ | PolyP polymers that can bind per activated platelet | - | Unknown |
| $k^*$ | Sensitivity of $k^{\text{cat}}$ to bound polyP concentration | - | Unknown |
| $K_D$ | Dissociation constant for polyP binding to APSs | - | Unknown |

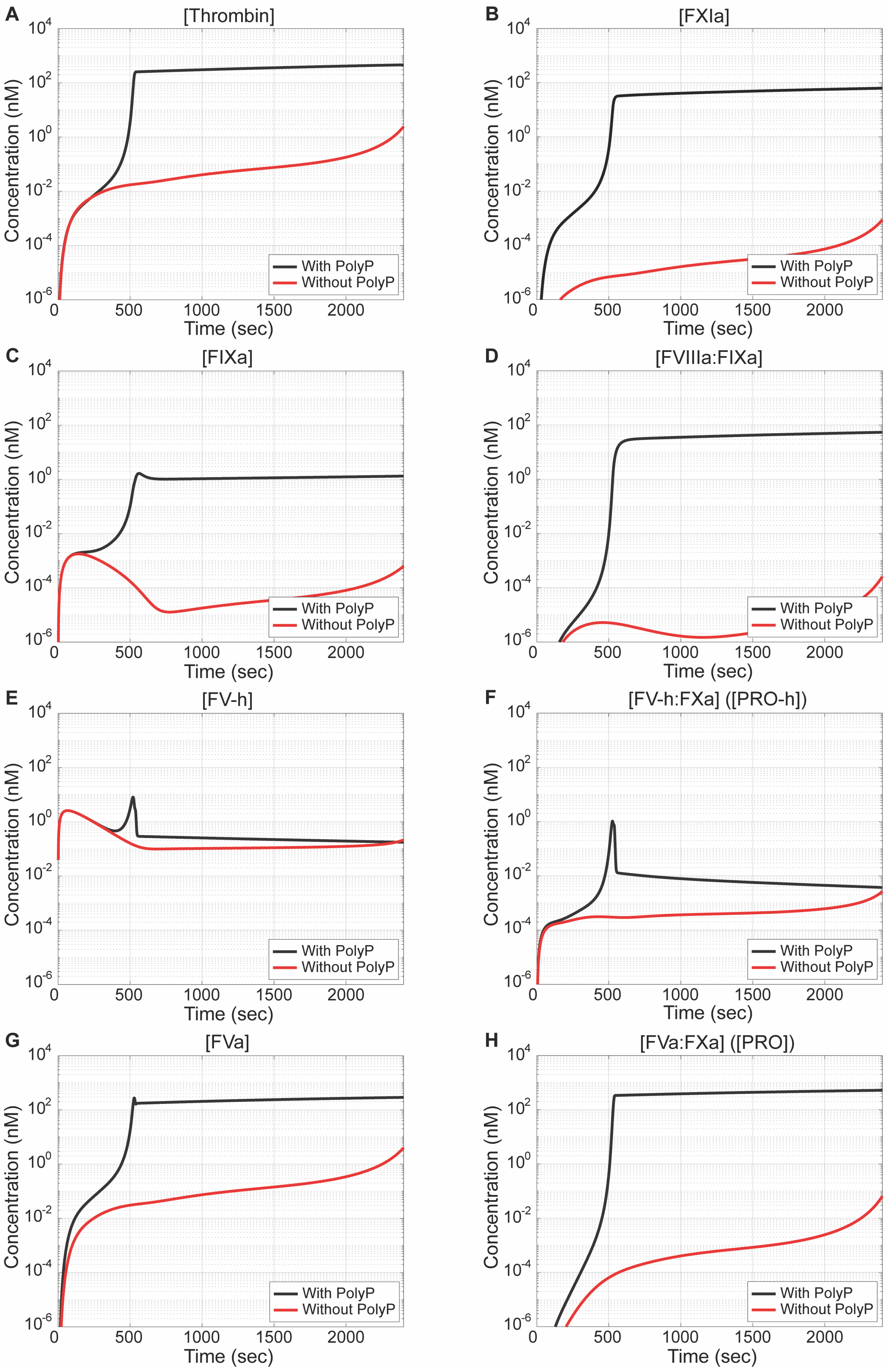


**Figure S1. PolyP enhances the speed of thrombin generation at low TF density in the implicit model. (Analog of entire Figure 2).** Time courses of enzyme concentrations with polyP (*black*) and without polyP (*red*): (A) Thrombin, (B) FXIa, (C) FIXa, (D) FVIIIa:FIXa (Tenase), (E) FVh, (F) FVh:FXa (PRO-h), (G) FVa, and (H) FVa:FXa (PRO). TF density = 0.3 fmol/cm$^2$; [TFPI] = 0.5 nM; shear rate = 100 s$^{-1}$.

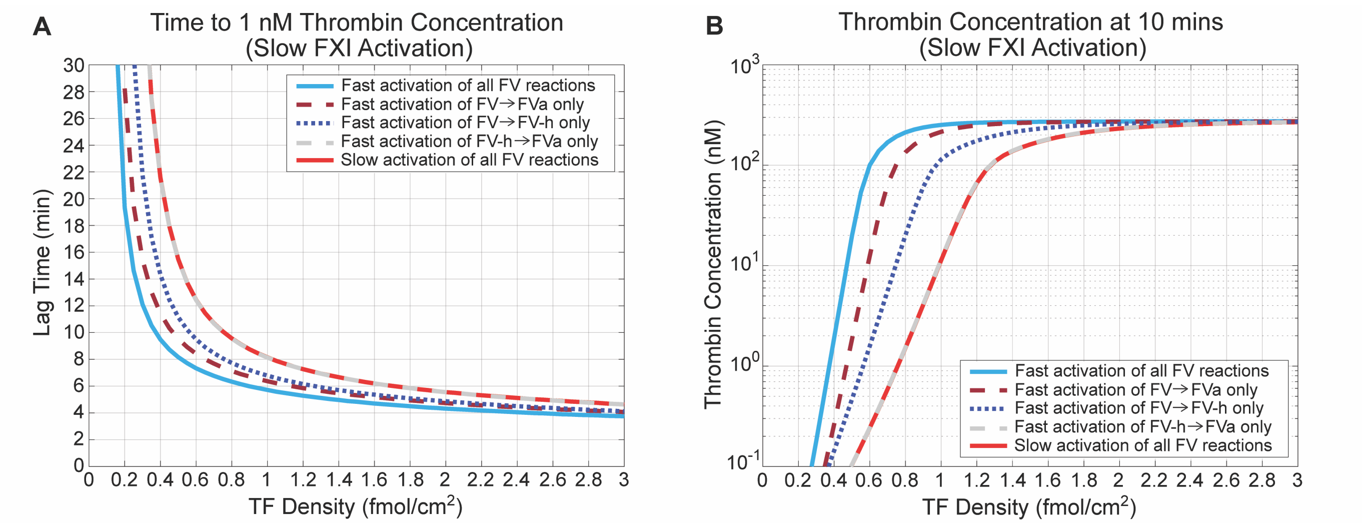


**Figure S2. Relative contributions of FV activation to thrombin generation with polyP under slow FXI activation in the implicit model. (Analog of Figure 4C-D).** (A) Time to 1 nM thrombin and (B) [thrombin] at 10 minutes with baseline FXI activation by thrombin and varying levels of acceleration of FV activation by FXa and thrombin. The curves corresponding to slow activation of all FV reactions (*solid red*) and to fast activation of FVh to FVa only (*dashed grey*) are on top of one another. "Fast" indicates that the respective reaction has been accelerated 10-fold for FV and 300-fold for FXI due to the presence of polyP, while "slow" indicates the baseline rate of activation without polyP. [TFPI] = 0.5 nM; shear rate = 100 $s^{-1}$.

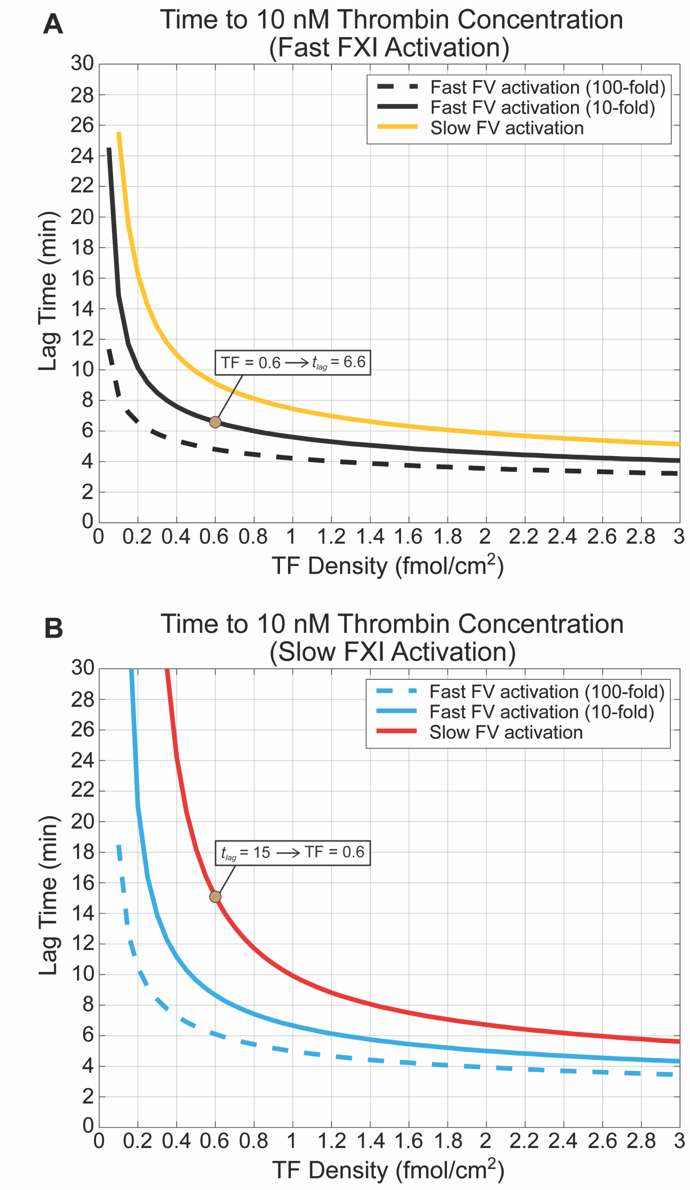


**Figure S3. The implicit model captures experimentally observed lag time to 10 nM thrombin concentration.** Time to 10 nM thrombin (A) with and (B) without accelerated FXI activation by thrombin. "Fast" indicates that the respective reaction has been accelerated due to the presence of polyP, while "slow" indicates the baseline rate of activation without polyP. 10-fold and 100-fold represents polyP-mediated enhancement of FV activation by FXa and thrombin. [TFPI] = 0.5 nM; shear rate = 100 $s^{-1}$. Given a TF density of 0.6 fmol/cm$^2$, times to 10 nM thrombin produced by the implicit model with slow FXI and FV activations (filled circle, panel B) and with fast FXI and FV activations (filled circle, panel A) roughly reproduce previously reported (Figure 3E of [7]) times to 10 nM thrombin in the absence and presence of polyP, respectively.

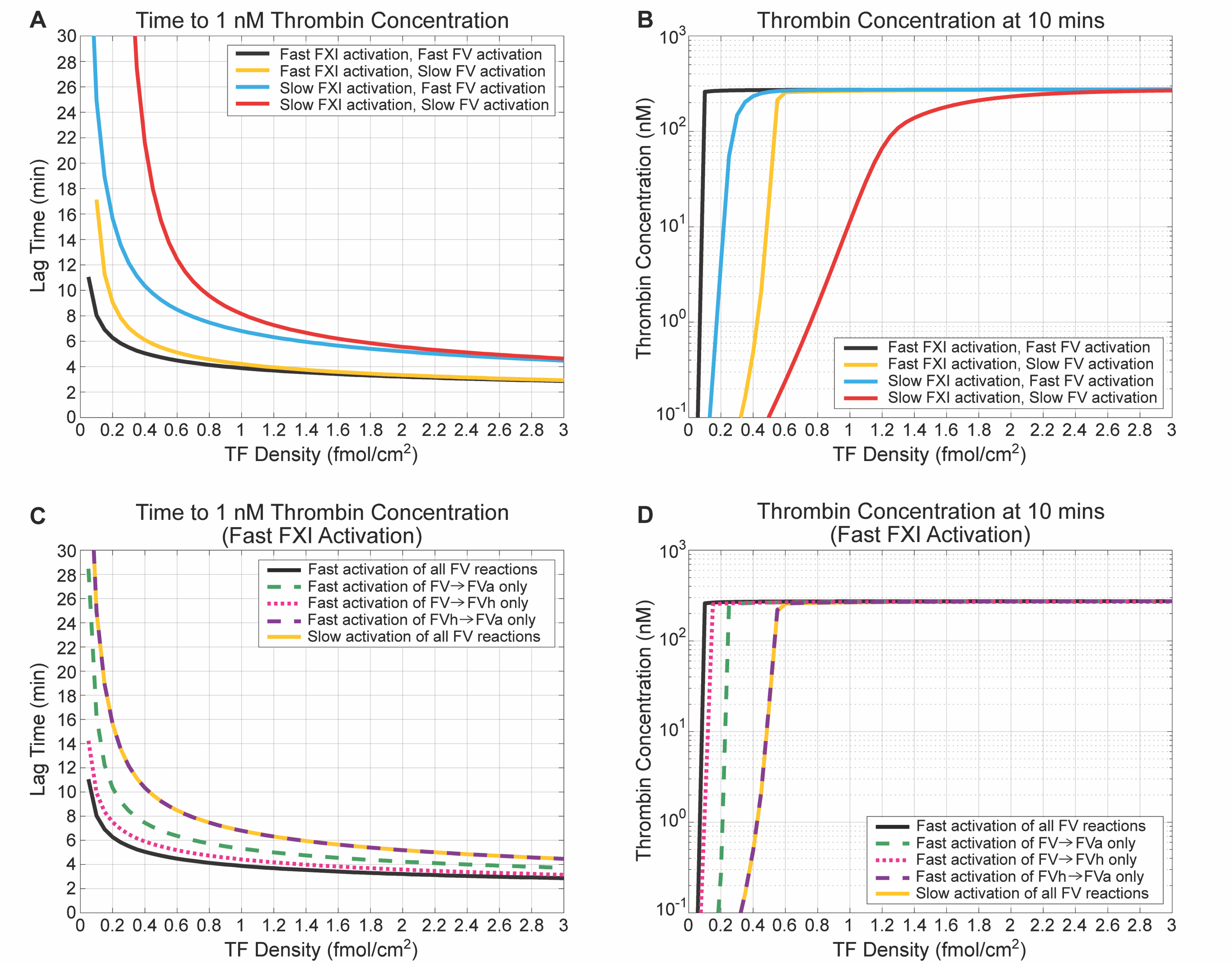


**Figure S4. Relative contributions of FXI and FV activation to thrombin generation with polyP under 100-fold enhancement of FV activation in the implicit model. (Analog of entire Figure 4).** (A) Time to 1 nM thrombin and (B) [thrombin] at 10 minutes with varying levels of acceleration of FXI activation by thrombin and FV activation by FXa and thrombin. (C) Time to 1 nM thrombin and (D) [thrombin] at 10 minutes with enhanced FXI activation by thrombin and varying levels of acceleration of FV activation by FXa and thrombin. The curves corresponding to slow activation of all FV reactions (*solid red*) and to fast activation of FVh to FVa only (*dashed grey*) are on top of one another. "Fast" indicates that the respective reaction has been accelerated 100-fold for FV and 300-fold for FXI due to the presence of polyP, while "slow" indicates the baseline rate of activation without polyP. With a 100-fold enhancement, acceleration of FV activation contributes more strongly to thrombin generation than accelerated FXI activation, but their combined effect is still sub-additive. Moreover, when FXI acceleration is accelerated, enhanced conversion of FV to FVh by FXa and FV to FVa by thrombin contribute comparably. The former reaction is dominant and produces similar results as all FV reactions accelerated, while the effect of accelerating FVh to FVa activation by thrombin remains negligible. [TFPI] = 0.5 nM; shear rate = 100 $s^{-1}$.

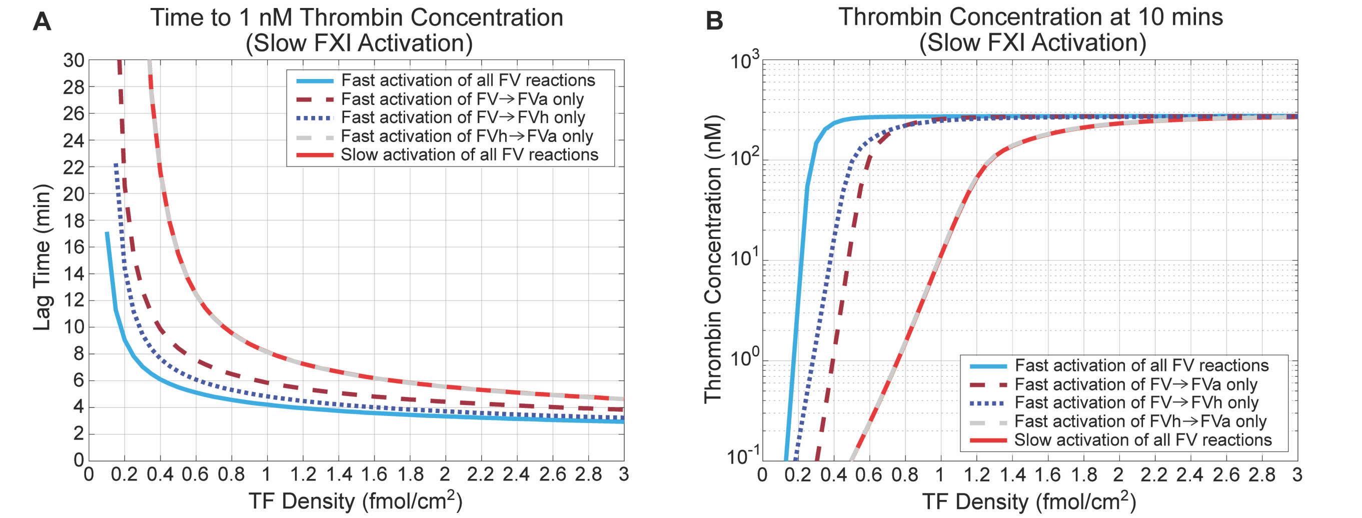


**Figure S5. Relative contributions of FV activation to thrombin generation with polyP under slow FXI activation and 100-fold enhancement of FV activation in the implicit model. (Analog of Figure 4C-D with two changes).** (A) Time to 1 nM thrombin and (B) [thrombin] at 10 minutes with baseline FXI activation by thrombin and varying levels of acceleration of FV activation by FXa and thrombin. The curves corresponding to slow activation of all FV reactions (*solid red*) and to fast activation of FVh to FVa only (*dashed grey*) are on top of one another. "Fast" indicates that the respective reaction has been accelerated 100-fold for FV and 300-fold for FXI due to the presence of polyP, while "slow" indicates the baseline rate of activation without polyP. With a 100-fold enhancement and baseline FXI activation, enhanced conversion of FV to FVh by FXa and FV to FVa by thrombin contribute comparably. The effect of accelerating FVh to FVa activation by thrombin remains negligible. [TFPI] = 0.5 nM; shear rate = 100 $s^{-1}$.

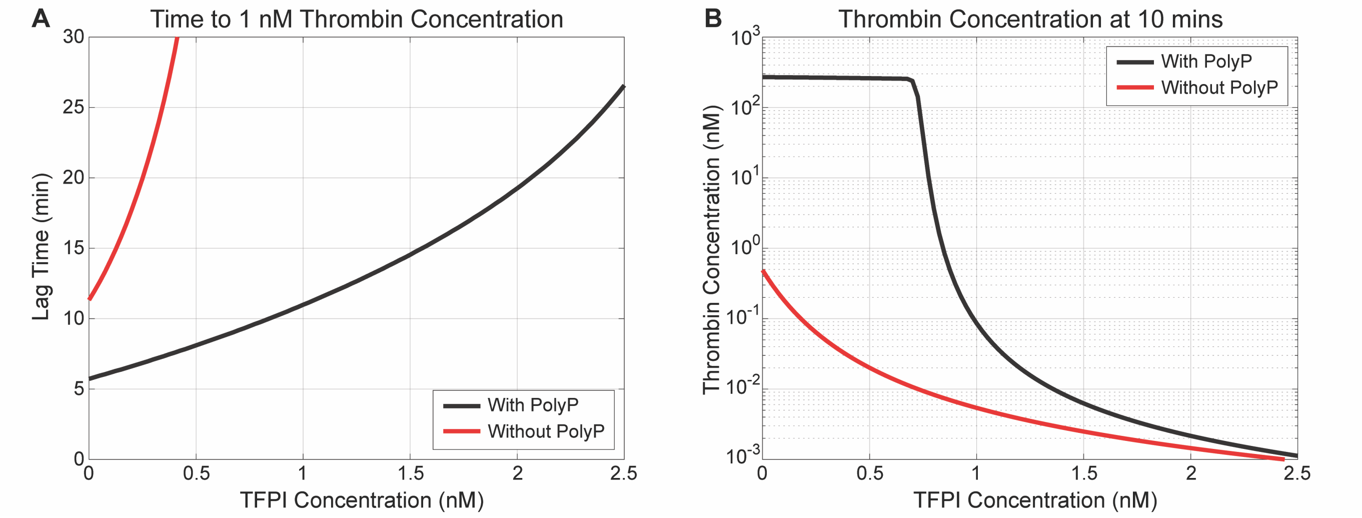


**Figure S6. PolyP reduces the sensitivity of thrombin generation to TFPI function at low TF density in the implicit model. (Analog of Figure 5A-B).** (A) Time to 1 nM thrombin and (B) [thrombin] at 10 minutes versus TFPI concentration with polyP (*black*) and without polyP (*red*). TF density = 0.3 fmol/$cm^2$; shear rate = 100 $s^{-1}$.

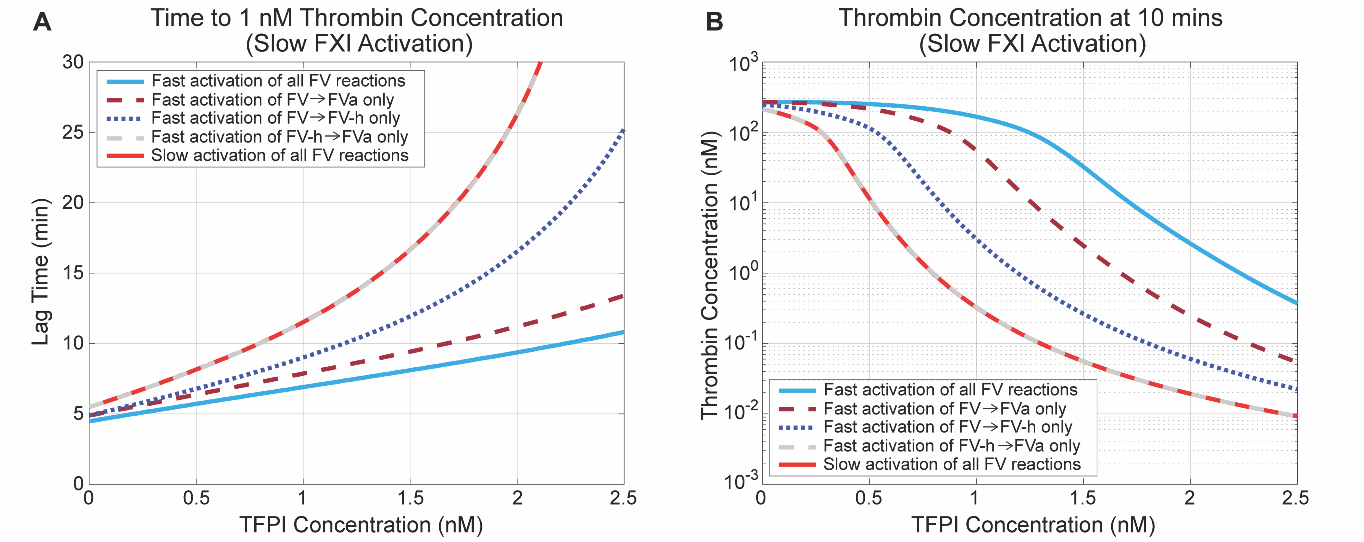


**Figure S7. Relative contributions of FV activation to sensitivity of thrombin generation to TFPI function under slow FXI activation in the implicit model. (Analog of Figure 5E-F).** (A) Time to 1 nM thrombin and (B) [thrombin] at 10 minutes versus TFPI concentration with baseline FXI activation by thrombin and varying levels of acceleration of FV activation by FXa and thrombin. The curves corresponding to slow activation of all FV reactions (*solid red*) and fast activation of FVh to FVa only (*dashed grey*) overlap. "Fast" indicates that the respective reaction is accelerated 10-fold for FV due to the presence of polyP, while "slow" indicates the baseline rate of activation without polyP. TF density = 1 fmol/cm$^2$; shear rate = 100 s$^{-1}$.

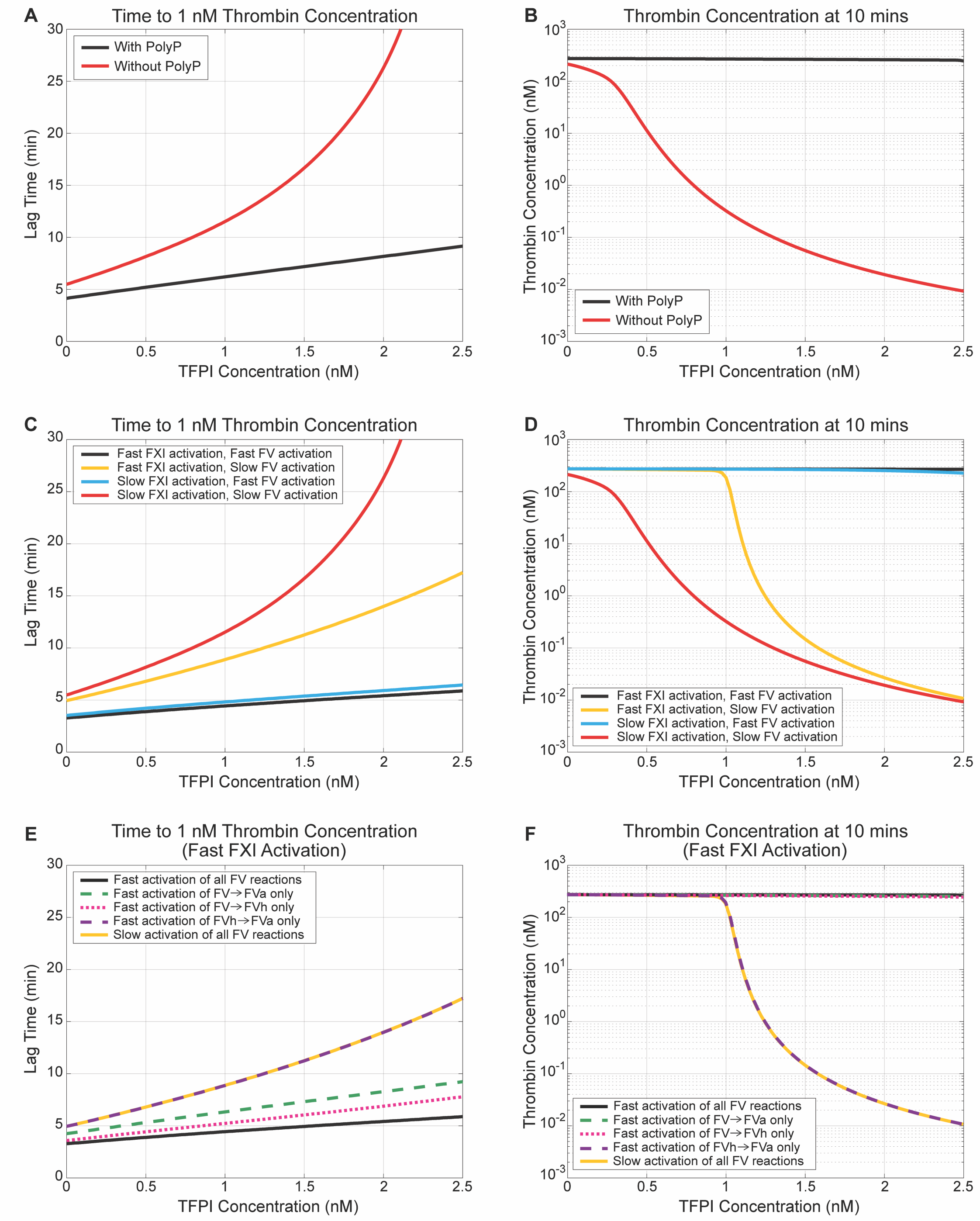


**Figure S8. PolyP decreases the sensitivity of thrombin generation to TFPI function under 100-fold enhancement of FV activation in the implicit model. (Analog of entire Figure 5).** (A-F) Time to 1 nM thrombin (*left*) and [thrombin] at 10 minutes (*right*) versus TFPI concentration: (A-B) with polyP (black) and without polyP (red), (C-D) with varying levels of acceleration of FXI activation by thrombin and FV activation by FXa and thrombin, and (E-F) with accelerated FXI activation by thrombin and varying levels of acceleration of FV activation by FXa and thrombin. In (E-F), the curves corresponding to slow activation of all FV reactions (*solid yellow*) and fast activation of FVh to FVa only (*dashed purple*) overlap. "Fast" indicates that the respective reaction is accelerated 100-fold for FV due to the presence of polyP, while "slow" indicates the baseline rate of activation without polyP. With a 100-fold enhancement, acceleration of FV activation contributes more strongly to reduced TFPI sensitivity to thrombin generation and lag time than accelerated FXI activation. Moreover, when FXI acceleration is accelerated, enhanced conversion of FV to FVh by FXa and FV to FVa by thrombin contribute comparably and result in high TFPI insensitivity. Both reactions produces similar results as all FV reactions accelerated, while the effect of accelerating FVh to FVa activation by thrombin remains negligible. TF density = 1 fmol/cm$^2$; shear rate = 100 s$^{-1}$.

| Average Sensitivity Measure<br>Slope of secant line of lag time from [TFPI] = 0 nM to [TFPI] = 1.0 nM (min/nM) | | |
|---|---|---|
| TF Density (fmol/cm$^2$) | Without PolyP | With PolyP |
| 0.1 | - | 17.9 |
| 0.3 | - | **5.3** |
| 0.5 | 29.2 | **3.4** |
| 0.7 | 11.6 | **2.6** |
| 0.9 | 7.1 | **2.2** |
| 1.0 | **6.0** | **2.1** |
| 1.5 | **3.6** | 1.6 |
| 2.0 | **2.7** | 1.3 |
| 2.5 | **2.3** | 1.2 |
| 3.0 | **1.9** | 1.1 |

**Figure S9. PolyP modulates the sensitivity of thrombin generation to TFPI in a TF-dependent manner, alternative sensitivity metric. (Analog of Figure 6E).** Average sensitivity measure (slope of secant line of lag time from 0 to 1 nM TFPI) with and without polyP for TF densities 0.1, 0.3, 0.5, 0.7, 0.9, 1, 1.5, 2, 2.5, and 3 fmol/cm$^2$. Bold entries indicate similar sensitivity slopes, occurring at lower TF with polyP and higher TF without polyP. A "-" in the table indicates that [thrombin] for the corresponding TF density never reached 1 nM for some TFPI concentrations between 0 and 1 nM. Shear rate = 100 s$^{-1}$.

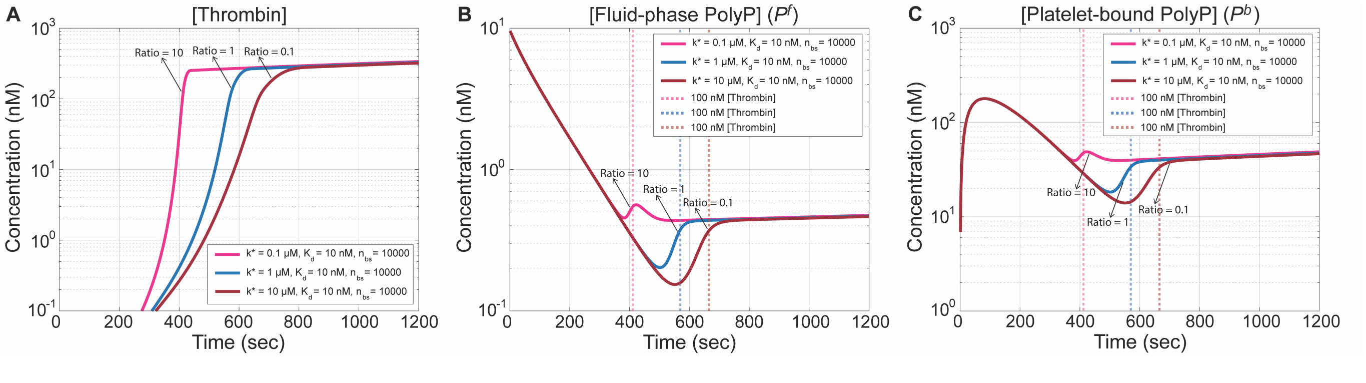


**Figure S10. Effect of the explicit model parameter ratio $R$ on thrombin and polyP concentrations.** Time courses of species concentrations: (A) Thrombin, (B) $P^f$, and (C) $P^b$ for $R = (n_{bs}/K_D)/k^*$ values of 0.1, 1, or 10 nM$^{-2}$. Note that the y-axis scales are different in each panel. Vertical dashed lines indicate the time to reach 100 nM [thrombin] in each case. Larger $R$ values lead to earlier and more rapid thrombin generation. The values of $R$ strongly influence polyP concentrations from ∼400-700 seconds, where the polyP curves for different values of $R$ separate (panels B and C). TF density = 1 fmol/cm$^2$; [TFPI] = 0.5 nM; shear rate = 100 s$^{-1}$.
